# Too dim, too bright, and just right: Systems analysis of the *Chlamydomonas* diurnal program upon acclimation to light stress and limitation

**DOI:** 10.1101/2024.10.22.618525

**Authors:** Sunnyjoy Dupuis, Valle Ojeda, Sean D. Gallaher, Samuel O. Purvine, Anne G. Glaesener, Raquel Ponce, Carrie D. Nicora, Kent Bloodsworth, Mary S. Lipton, Krishna K. Niyogi, Masakazu Iwai, Sabeeha S. Merchant

**Affiliations:** Department of Plant and Microbial Biology, University of California, Berkeley, CA 94720, USA; California Institute for Quantitative Biosciences, University of California, Berkeley, CA 94720, USA; Earth and Biological Sciences Division, Pacific Northwest National Laboratory, Richland, WA 99352, USA; Howard Hughes Medical Institute, University of California, Berkeley, CA 94720, USA; Molecular Biophysics and Integrated Bioimaging Division, Lawrence Berkeley National Laboratory, Berkeley, CA 94720, USA; Environmental Genomics and Systems Biology Division, Lawrence Berkeley National Laboratory, Berkeley, CA 94720, USA; Department of Molecular and Cell Biology, University of California, Berkeley, CA 94720, USA

**Keywords:** photoacclimation, RNA-Seq, proteomics, lipidomics, pigments, chloroplast, microscopy, NPQ, alga, chlorophyte

## Abstract

Photosynthetic organisms coordinate their metabolism and growth with diurnal light, which can range in intensity from limiting to inhibitory. To gain a comprehensive understanding of how diurnal regulatory circuits interface with sensing and response to various light intensities, we performed a systems analysis of synchronized *Chlamydomonas* populations acclimated to low, moderate, and high diurnal light. Transcriptomic and proteomic data revealed that the *Chlamydomonas* rhythmic gene expression program is resilient to limiting and excess light. Although gene expression and photodamage are dynamic over the diurnal cycle, *Chlamydomonas* populations acclimated to low and high diurnal light exhibit constitutive phenotypes with respect to photosystem abundance, thylakoid architecture, and non-photochemical quenching that persist through the night. This suggests that cells “remember” or anticipate the daylight environment. The integrated data constitute an excellent resource for understanding gene regulatory mechanisms and photoprotection in eukaryotes under environmentally relevant conditions.

## INTRODUCTION

Photosynthetic organisms have evolved under a highly dynamic light environment, where the intensity of solar irradiance varies over several orders of magnitude each day^1^. The periodic availability of sunlight over diurnal cycles restricts photosynthetic activity to the daytime, yielding daily rhythms in cellular growth and metabolic activity^2–4^. Over each day period, incident light intensity can range from limiting to excess. Under excess light, the rate of light absorption exceeds the rate of photosynthetic electron transfer, leading to production of reactive chemical species that can cause photooxidative damage. Accordingly, these organisms have evolved sophisticated mechanisms to sense light intensity, adjust light-harvesting capacities, dissipate excess absorbed energy, and repair photodamage^5–7^. In photosynthetic eukaryotes, these mechanisms are coordinated across cellular compartments^8^.

The unicellular, eukaryotic green alga *Chlamydomonas reinhardtii* (*Chlamydomonas* hereafter) is a classic model organism for studying photosynthesis and its regulation^9^. As a unicellular organism, the tissue- and developmental-stage-specific heterogeneities that complicate studies in land plants are avoided. Yet, having a shared evolutionary history with land plants, *Chlamydomonas* harbors conserved photosynthetic machinery, regulatory pathways, and photoprotective mechanisms. When the amount of absorbed light exceeds the capacity for photochemical quenching by charge separation, non-photochemical quenching (NPQ) mechanisms are induced that dissipate energy as heat. These mechanisms operate over both short and long timescales^10,11^. The most rapidly reversible NPQ, called energy-dependent quenching or qE, is performed by energy-dissipative light-harvesting complex (LHC) proteins (e.g., LHCSRs) that accumulate in thylakoid membranes alongside the photosynthetic apparatus in high light (HL)^12–16^. In addition, a slowly reversible NPQ, called zeaxanthin-dependent quenching or qZ, occurs when high rates of linear electron transfer from photosystem II (PSII) to photosystem I (PSI) increase the proton gradient across the thylakoid membrane (ΔpH), activating violaxanthin de-epoxidase (CVDE1) which converts violaxanthin (Vio) to zeaxanthin (Zea), an energy-dissipative carotenoid^17–19^. Another slowly reversible NPQ, called state-transition-dependent quenching or qT, occurs when the high reduction state of the plastoquinone pool activates the STT7 kinase, which phosphorylates the LHC proteins of PSII (LHCII)^20^. Phosphorylated LHCII trimers associate with PSI in State 2, thereby redistributing excitation energy between the photosystems^20,21^. Upon prolonged HL exposure, photoinhibition itself can cause quenching in the PSII reaction center, known as qI, the most slowly reversible NPQ mechanism^22^.

In addition to NPQ, *Chlamydomonas* can acclimate to prolonged HL by decreasing chlorophyll content and antenna size to reduce the excitation energy pressure on the photosystems, a process termed photoacclimation^23–32^. Most studies on photoacclimation in *Chlamydomonas* have used cultures maintained in continuous light to maximize growth rate or cultures shifted from one continuous light intensity to another^15,23,28,30,31,33–38^. However, in natural environments, photoacclimation occurs in the context of the diurnal light cycle, over which gene expression, metabolism, and physiology are coordinated with the time of day. Cell growth (G1 phase of the cell cycle) is restricted to the light phase. After a full day of growth, *Chlamydomonas* cells replicate their genomes (S phase) and undergo mitotic divisions (M phase) to produce daughter cells of equal sizes^39^. As a result, *Chlamydomonas* populations synchronize when grown under repeated diurnal cycles in the laboratory.

With millions of cells each at the same stage of the cell cycle, synchronous populations produce high signal-to-noise for measuring changes in gene expression and metabolism that would be obscured in batch cultures. Synchronous *Chlamydomonas* populations have been a workhorse for studying the cell cycle^40^. They have also revealed that metabolic pathways and the biogenesis of various cellular structures (e.g., photosynthetic and respiratory complexes, cilia, ribosomes) are segregated to specific times of day to optimize resource allocation over the diurnal cycle. Our previous work on synchronous populations grown under moderate diurnal light (ML) revealed that this diurnal program is achieved through rhythmic accumulation of ∼85% of nuclear and organellar transcripts^41^. Interestingly, *LHCSRs* were among these rhythmically expressed genes, even at low light (LL), suggesting that the photoprotective mechanisms that *Chlamydomonas* engages could also be dynamic over the diurnal cycle. Only a few studies to date have explored this hypothesis. Nawrocki *et al.* (2020) found that under sinusoidal HL illumination, NPQ capacity remained high over time, which was attributed to the maintenance of LHCSR proteins during light and dark phases^42^. Using a similar sinusoidal light regime, van den Berg and Croce (2022) found that carotenoid composition varied significantly over the diurnal cycle in *Chlamydomonas*^43^.

Apart from these findings, it is largely unclear how diurnal rhythms in gene expression and metabolism influence photoprotection, and conversely, how the intensity of light may impact *Chlamydomonas’* diurnal program. As diurnal illumination is a fundamental characteristic of our planet, addressing these questions is critical to our understanding of photosynthetic organisms, their capacity to acclimate to changing environments, and our ability to engineer them for improved productivity. To understand how diurnal regulatory circuits interface with the sensing and response to different light intensities, we performed a systems analysis of synchronized *Chlamydomonas* populations acclimated to diurnal LL, ML, and HL. We present a comprehensive view of gene expression, photosynthesis, metabolism, and chloroplast morphology at five timepoints across the day and night that integrates transcriptomics, proteomics, lipidomics, pigments, bioimaging, and biophysical data. We find that diurnal rhythms in gene expression are resilient to limiting and excess light, but that diurnal photoacclimation gives rise to hundreds of gene expression changes, even at night. Although gene expression is highly dynamic, we found that the acclimated populations maintained altered thylakoid architecture and photosystem abundance into the night phase. In this way, *Chlamydomonas* may prepare for the light environment of a typical day.

## RESULTS

### The *Chlamydomonas* diurnal program is resilient to challenging light intensities

To understand how light intensity influences the *Chlamydomonas* diurnal program, we acclimated synchronous populations to one of three intensities of diurnal light and then performed analytical measurements over a diurnal cycle time course. The populations were grown photoautotrophically in turbidostat photobioreactors for >2 weeks under 12-h-dark/12-h-light cycles of either LL, ML, or HL (∼50, ∼200, or ∼1000 µmol photons m^−2^ s^−1^, respectively). The ML condition, which produces an exquisite level of synchrony in which cells divide once per diurnal cycle upon nightfall, has been studied extensively and served as our control^41^. We assayed physiology and gene expression of the acclimated populations at five timepoints, numbered relative to the dark-to-light transition: two hours before the light phase (–2), two, six, and ten hours into the light phase (+2, +6, +10), and two hours into the dark phase, which is ten hours before the next light phase (–10) (Figure 1A). Thus, with three photoacclimated populations and five timepoints, we compared a total of 15 cellular states (Figure 1B).

**Figure 1:**
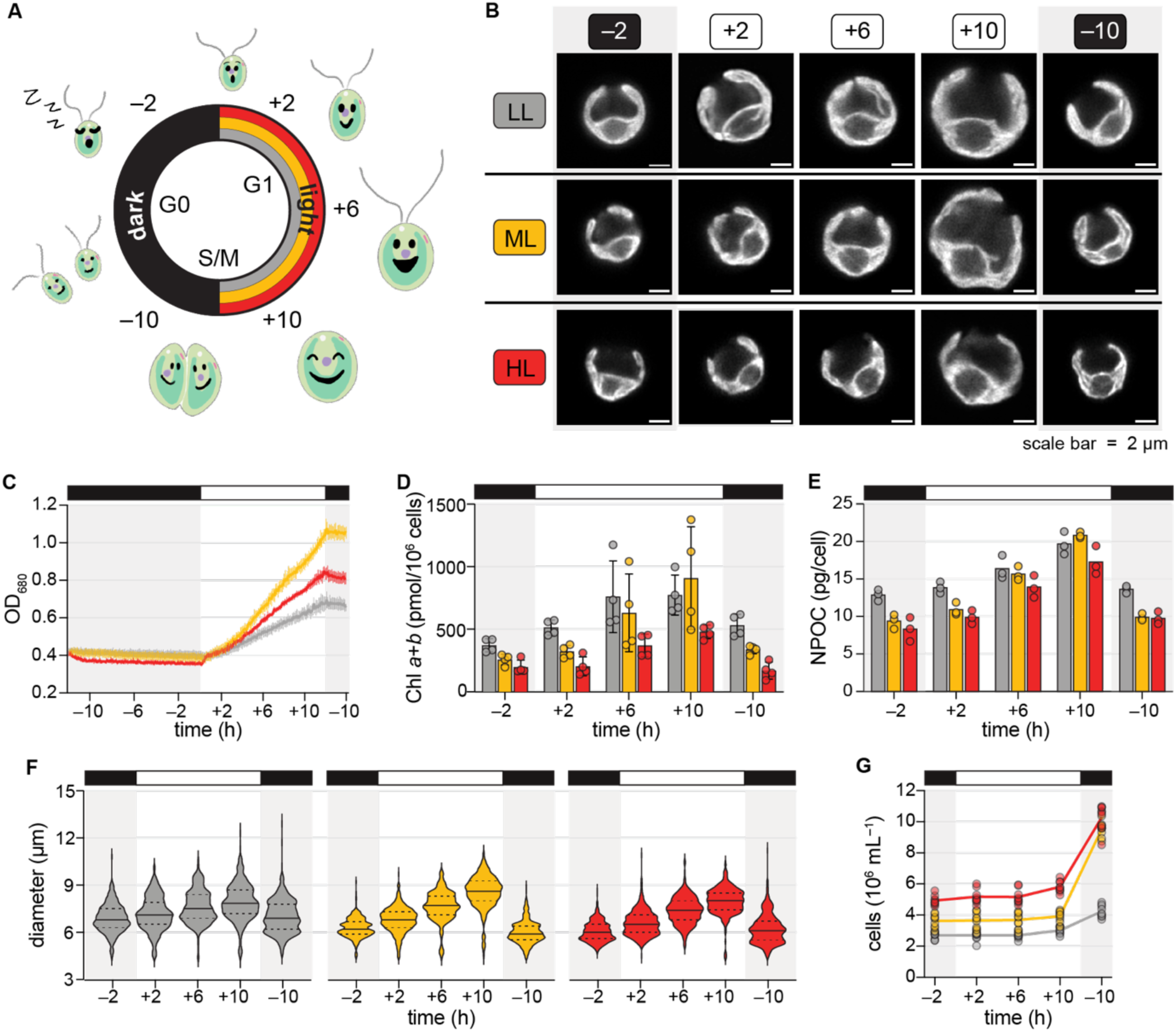
The *Chlamydomonas* cell cycle is coordinated with the time of day even under challenging light intensities. (A) Schematic of *Chlamydomonas* populations synchronized under 12-h-dark/12-h-light cycles of LL (grey), ML (yellow), and HL (red). (B) Representative live-cell Chl fluorescence images of the 15 cellular states assayed in this study. (C) Optical density (OD_680_) of all cultures was set to 0.4 at the beginning of the dark phase and continuously monitored over 26 h. Data are represented as the mean of 9 experimental replicates (n = 9) with error bars representing the standard deviation. (D) Cellular Chl content. (E) Cellular organic carbon content, estimated as non-purgeable organic carbon (NPOC). (F) Cell diameter; solid lines represent the median and dashed lines the quartiles of 100 cells from each of 3 experimental replicates (n = 300). (G) Cell density.

The acclimation conditions yielded excellent reproducibility of culture growth across independent replicate experiments performed months apart (Figure 1C). Regardless of light intensity, growth was restricted to the light phase. The LL-acclimated population grew the most slowly and reached the lowest final density, indicating that light energy input limits growth in LL. The HL-acclimated population also reached a lower density than the ML population did (*p* < 0.05), suggesting that this light intensity causes photoinhibition. As expected, LL-acclimated cells had more Chl compared to HL-acclimated cells (Figure 1D, *p* < 0.05). Interestingly, this difference was maintained even after 10 h of dark (–2). On average, LL-acclimated cells also had a higher organic carbon content than did the ML and HL populations at the –2, +2, and –10 timepoints (Figure 1E, *p* < 0.05). This reflected a significant increase in the average size of LL cells at these timepoints (Figure 1F, *p* < 0.05), as fewer LL cells were able to divide upon the light-to-dark transition. The number of cells in the ML and HL populations increased over two-fold on average, whereas the LL population did not quite double (mean fold-change = 1.58) (Figure 1G). Nonetheless, cell division was coordinated with the light-to-dark transition in all three populations.

To understand how acclimation to different light intensities impacts *Chlamydomonas’* rhythmic gene expression program, we performed RNA-Seq and tandem mass tag (TMT) proteomics (Tables S1 and S2). We detected 16,735 nucleus-encoded mRNAs (95%) (Figure S1A), 10,011 nucleus-encoded proteins (56.9%), 59 chloroplast-encoded proteins (81.9%), and 5 mitochondria-encoded proteins (62.5%) (Figure S1B). Genome-wide, mRNA and protein abundances appeared to oscillate with a 24 h period over the diurnal cycle, and light intensity seemed to have little effect on the timing of accumulation. Principal component analysis (PCA) of the RNA-Seq data confirmed that most of the variation in the transcriptome is explained by time of day, rather than by light intensity, and it also confirmed tight reproducibility across experimental replicates collected weeks apart (Figure 2A). PCA of the TMT proteomics data showed that time of day also drives the variation in protein abundances (Figures 2B).

**Figure 2:**
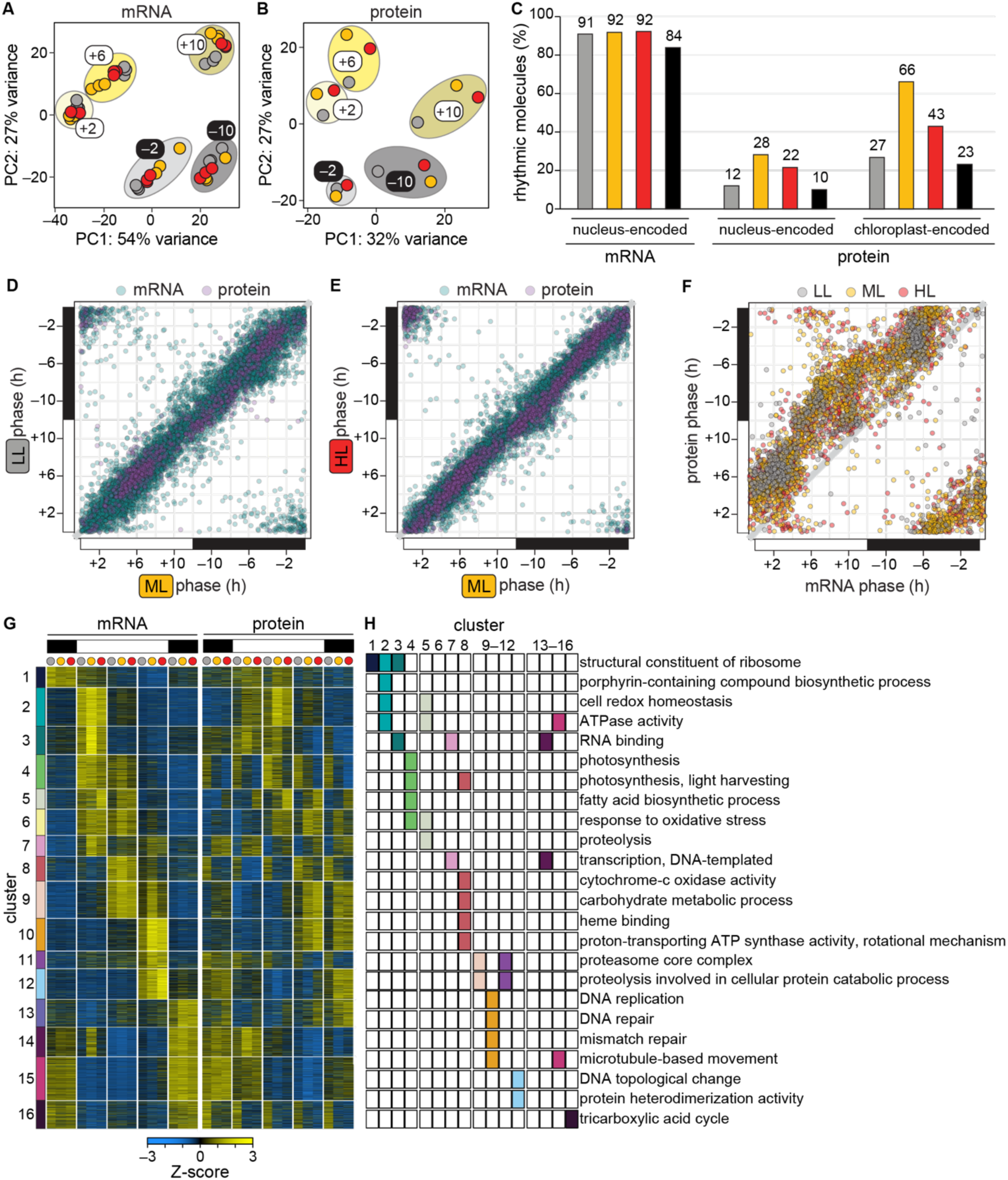
The rhythmic diurnal gene expression program is maintained under diurnal LL and HL. (A) PCA of the transcriptome; ellipses demonstrate grouping by time of day. (B) PCA of the mean proteome; ellipses demonstrate grouping by time of day. (C) Proportion of detected molecules that exhibit a significant diurnal rhythm in abundance in diurnal LL (grey), ML (yellow) and HL (red) and in all three light intensities (black). No significant rhythms were detected for the 5 mitochondria-encoded proteins captured by TMT proteomics. (D) Correlation between the phase of rhythmic gene expression in diurnal LL and ML. (E) Correlation between the phase of rhythmic gene expression in diurnal HL and ML. (F) Correlation between the phase of expression for rhythmic mRNAs and their cognate proteins; grey line represents a 1:1 correlation. (G) Normalized mRNA and protein levels for 9,821 genes grouped into 16 clusters. (H) Selected GO terms significantly enriched in the 16 clusters in Figure 2G. See also Figure S1 and Tables S1, S2, S3, and S4.

Next, we used the algorithm DiscoRhythm to determine which genes were expressed with a diurnal periodicity in the three populations and to estimate the phase of these rhythms (the time at which expression peaks)^44,45^ (Table S3). We found that roughly 91% of nucleus-encoded transcripts exhibited a significant diurnal oscillation in the populations, whereas significant diurnal rhythms were only detected for 12–28% of nucleus-encoded proteins depending on the diurnal light intensity (Figure 2C). A similar study which instead used data-independent acquisition proteomics in the marine alga *Ostreococcus* also found decreased rhythmicity of the proteome relative to the transcriptome over diurnal cycles: 80% of transcripts and 55% of proteins exhibited a diurnal rhythm^46^. However, as both PCA and *k*-means clustering suggested that a large proportion of the *Chlamydomonas* proteome was rhythmically expressed (Figures 2B and S1B), DiscoRhythm may underestimate rhythmicity in TMT proteomics data due to the compressed dynamic range. On the other hand, 26–66% of chloroplast-encoded proteins accumulated with a significant rhythm.

For rhythmic mRNAs and proteins alike, the phase of expression in the LL and HL populations correlated well with the phase in the ML control population (Figures 2D and 2E). Thus, the timing of gene expression is maintained regardless of the intensity of diurnal light in *Chlamydomonas*. As predicted by the central dogma of biology (DNA to RNA to protein), we found that the phase for rhythmically expressed proteins often fell 2-to-8 hours after the phase for the cognate mRNA (Figure 2F).

Both mRNA and protein products were detected from 9,821 genes, and these genes were grouped into clusters based on their expression patterns (Figure 2G). 14 of the 16 clusters were significantly enriched for gene ontology (GO) terms, revealing that the expression of genes encoding related functions was gated to specific times of day (as reported previously^41,47^) and hinting at light-responsive changes in cellular physiology (Figures 2H and S1C, Table S4). For example, Cluster 4 genes – enriched for “photosynthesis, light harvesting,” “fatty acid biosynthesis,” and “response to oxidative stress” terms – peaked in mRNA abundance early and midday (+2 and +6), and the corresponding proteins were more abundant in the LL population at all timepoints. Cluster 5 mRNAs accumulated similarly to those in Cluster 4, but proteins in this cluster were instead more abundant in the HL population, especially at midday (+6). This cluster was also significantly enriched with “response to oxidative stress” in addition to proteolysis-related terms, suggesting that the HL population may experience photooxidative damage at midday. Cluster 10, enriched for DNA replication and repair terms, was most highly expressed at the end of the day (+10) when the cells were in S and M phase of the cell cycle. The magnitude of expression of these genes was reduced in the LL population, in line with the observation that fewer LL cells divided upon nightfall (Figure 1G).

Taken together, our results reveal that the *Chlamydomonas* diurnal program is resilient to light stress and limitation. Populations are relatively synchronized under diurnal HL and LL, making this an excellent system to investigate how *Chlamydomonas* copes with challenging light intensities at each phase of the diurnal cycle.

### Hundreds of genes are differentially expressed in populations acclimated to diurnal LL and HL, even during the night

Although the diurnal program was largely maintained, light intensity did alter the expression of many genes (Figures S1 and 2). Therefore, we performed differential expression analysis on the full *Chlamydomonas* transcriptome and proteome at each timepoint (Table S5). Hundreds of genes were differentially expressed relative to the ML control population at both the mRNA and protein levels, even at the –2 timepoint when all three populations had been in the dark for 10 h (Figures 3A and 3B). Thus, acclimation to different intensities of diurnal light influences *Chlamydomonas’* biology in both the daytime and the nighttime. We found that more mRNAs were differentially expressed in the LL population than in the HL population, which at least partially reflects the decreased synchrony of the LL population (Figure 3A). For both the LL and HL populations, the greatest number of changes was observed at the +2 timepoint. Dawn and the onset of excess light are both known to induce massive transcriptional changes in *Chlamydomonas*^36^ and to cause significant changes in mRNA half-life^48–51^. Our results show that even after prolonged diurnal photoacclimation, light intensity has the greatest impact on the transcriptome at the start of each day.

**Figure 3:**
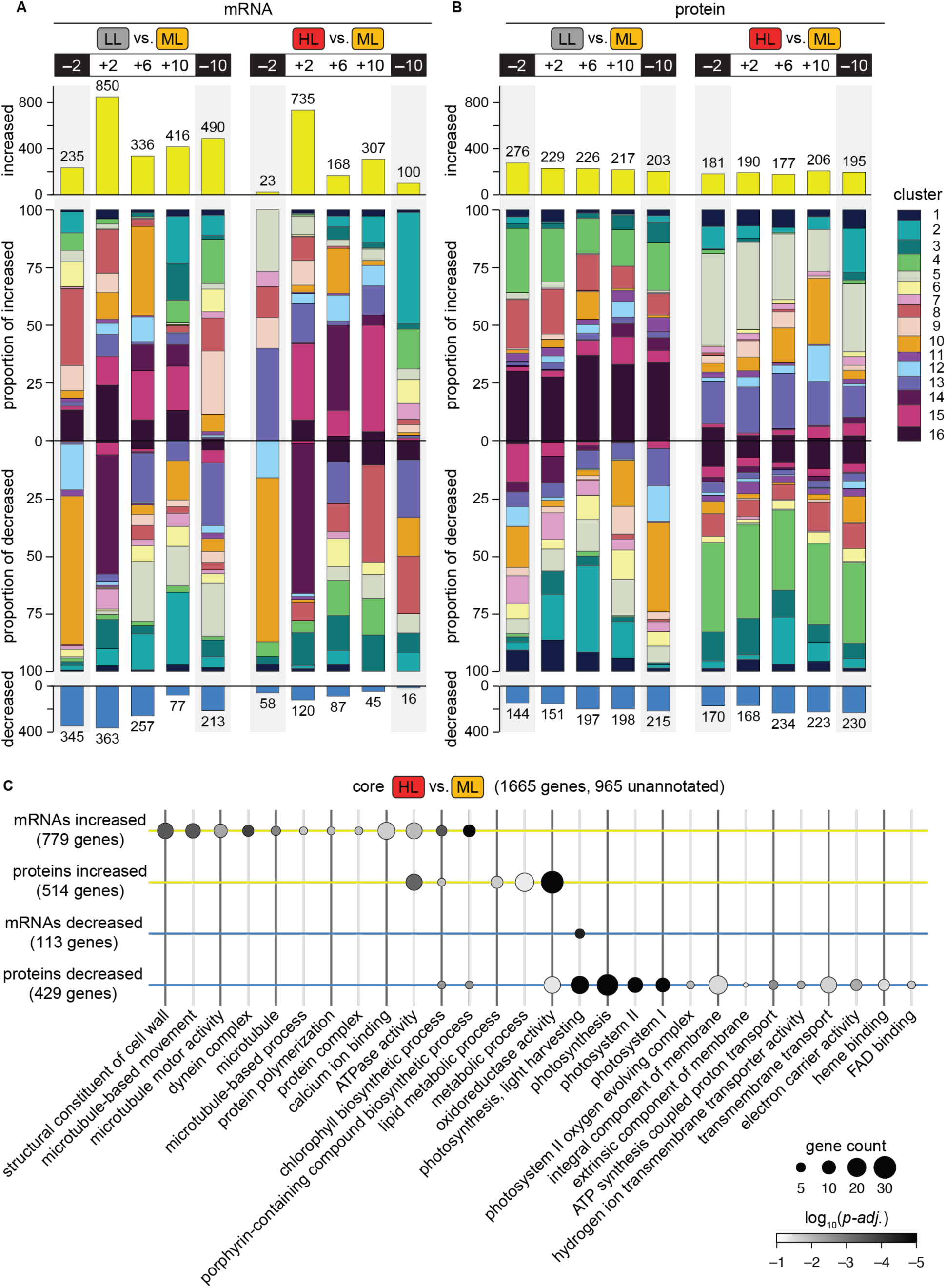
Hundreds of genes are differentially expressed in populations acclimated to diurnal LL and HL at all timepoints, and comparison reveals HL-specific changes related to motility and metabolism. (A) The number of mRNAs significantly increased (top) and decreased (bottom) in abundance in each comparison at each timepoint, and the proportion of those mRNAs that belong to each cluster in Figure 2G (center); the key at right is scaled by the proportion of total genes in each cluster for comparison. (B) Significant differences in protein abundance represented as in (A). (C) GO term enrichment in the “core HL-responsive” gene expression changes. See also Figures S2, S3, and S4, and Tables S5 and S6.

On the other hand, the number of changes to the proteome was quite similar at each timepoint (Figure 3B). In addition, proteins from the same clusters were often increased or decreased at all five timepoints, suggesting that diurnal photoacclimation results in constitutive changes in protein abundance across the diurnal cycle. Comparative analysis revealed that a substantial proportion of the changes to the proteome were constitutive: 13% of the proteomic responses to diurnal LL and 24% of the proteomic responses to HL occurred at all five timepoints (Figure S2). In contrast, only 1% of mRNA changes were maintained across all five timepoints. This difference likely reflects the increased rhythmicity of the transcriptome relative to the proteome (Figure 2C) and inherent differences in the half-lives of mRNAs and proteins.

We noted that differentially expressed mRNAs often belonged to a different set of clusters than did the differentially expressed proteins at a particular time (Figures 3A and 3B), suggesting that changes in the transcriptome often did not yield a corresponding change in the proteome. We determined the number of genes that were differentially expressed at both the mRNA and protein levels simultaneously or after a 4 h delay, as we had found that rhythmic proteins peaked later than their cognate transcripts (Figure 2F). Indeed, we found that most changes occurred only at the mRNA or protein level, even when we considered a 4 h delay (Figure S3). While methodological differences undoubtedly contribute to this lack of overlap, these results also speak to characteristic differences in the roles of mRNA and protein abundance in the cell and how they are controlled^52–55^.

We sought to generate a core list of HL-responsive genes from our data. We reasoned that gene expression changes which occurred in both the HL and LL populations likely reflect differences in productivity or synchrony relative to the ML control population rather than light-specific responses. Therefore, we defined core HL-responsive mRNAs and proteins as either 1) significantly increased in HL and unchanged or decreased in LL, or 2) significantly decreased in HL and unchanged or increased in LL. These criteria filtered out hundreds of gene expression changes, especially at the +2 timepoint: of the 735 mRNAs that were significantly increased at this time, 453 were specific to the HL-acclimated population (Figure S4). In total, 1665 genes had HL-specific responses in transcript or protein abundance at some point over the diurnal cycle – roughly 10% of the *Chlamydomonas* genome. The gene expression changes include expected responses such as increases in the expression of light-stress and light-inducible genes (e.g., *LHCSRs*, *ELIPs*) and decreases in photosystems and LHCs (Figure 3C, Table S6). In addition, CiliaCut genes – genes retained only in ciliated organisms^56^ – were significantly enriched in the core HL-responsive list (Figure S4B, *p* = 3.3 x10^−15^), and many microtubule- and dynein-related genes showed an HL-specific increase in transcript abundance, suggesting that diurnal HL triggers a motility response in *Chlamydomonas*^57,58^. Over half of the core HL-responsive genes (965 genes) are not associated with a functional annotation, representing a large space for discovery of genes important for acclimation to light stress under environmentally relevant conditions.

### Carbon fixation appears to be limited by CO_2_ availability in *Chlamydomonas* cells acclimated to diurnal HL

Having determined that the expression of genes related to photosynthesis and respiration was altered in the LL and HL populations (Figures 2 and 3), we next monitored respiratory and photosynthetic activity. We found that the rate of O_2_ consumption during dark incubation was lowest for cells collected during the night phase in all three populations, as observed previously in diurnal ML^41^ (Figure 4A, *p-adj.* < 0.05). The maximum rate of O_2_ evolution when provided light was also low for cells collected during the night phase (Figure 4B, *p-adj.* < 0.05). These temporal differences not only reflect how the cellular demand for ATP changes over the cell cycle (lower in G0) but also how the environment influences metabolic rate. Cooler temperatures during the night and changing CO_2_ affinity^59^ likely shape the respiratory and photosynthetic activity of the cells. Respiratory activity increased during the day but was significantly lower in LL cells (Figure 4A, *p* < 0.05). There was no significant difference in O_2_-evolution rate per biomass across the three populations at any timepoint (Figure 4B). However, when normalized to Chl, O_2_-evolution rate was similar for HL and ML cells and lower for LL cells (Figure 4C). Thus, LL cells had decreased metabolic activity despite having increased Chl content (Figure 1D) and the abundance of proteins enriched for photosynthesis and respiration GO terms (Figure 3B).

**Figure 4:**
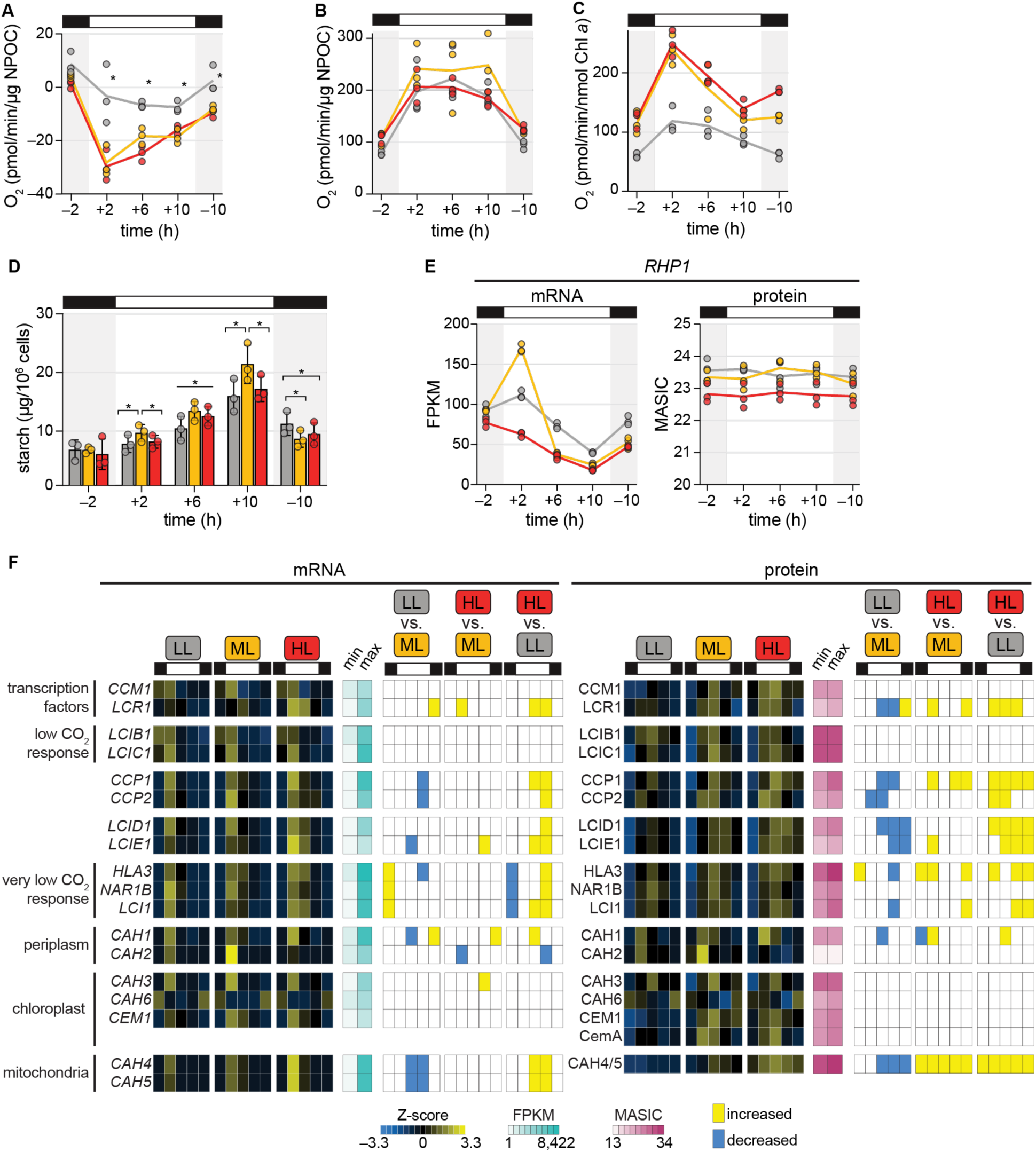
Carbon fixation appears to be limited by CO_2_ availability in *Chlamydomonas* cells acclimated to diurnal HL. (A) Rate of O_2_ consumption by cells collected at the designated timepoints and assayed in the dark for 5 min, normalized to NPOC as a proxy for biomass. (B) Rate of O_2_ evolution by cells collected at the designated timepoints and assayed at 1000 µmol photons m^−2^ s^−1^ for 3 min with excess CO_2_, normalized to NPOC as a proxy for biomass. (C) Rate of O_2_ evolution measured as in (B) but normalized to mean Chl *a* content. (D) Cellular starch content. (E) *RHP1* gene expression as a proxy for relative intracellular CO_2_ concentration. (F) Changes in mRNA and protein abundance of genes related to intracellular C_i_ concentration.

A fraction of fixed carbon and reductant derived from photosynthesis is stored as starch, which is synthesized during the day and then degraded at night to sustain metabolism in the dark. Starch metabolism is subject to redox regulation and is under circadian control in plants and *Chlamydomonas*^41,60–62^. Previous work has shown that starch accumulation increases upon an increase in light intensity^36^, serving as a sink for excess reductant, so we hypothesized that starch content would be highest in the HL population. However, we found that *Chlamydomonas* cells accumulated significantly less starch over the light phase in both LL and HL (Figure 4D, *p* < 0.05). Cellular starch content decreased to an equivalent level in the three populations by the end of the night (–2). Starch degradation rate is known to be adjusted to the anticipated length of the night phase in the land plant *Arabidopsis thaliana*^63^, and our results suggest that the rate of starch degradation during the night may be similarly adjusted in *Chlamydomonas* to achieve a set starch content before dawn.

Since HL cells and ML cells exhibited similar maximum O_2_-evolution rates, we hypothesized that CO_2_ concentration limits carbon fixation and starch synthesis in our HL condition. To explore this hypothesis, we investigated the expression of the CO_2_ channel RHP1, a marker for intracellular CO_2_ levels which increases at both the transcript and protein level in high CO_2_^64,16^. We found that *RHP1* transcript levels were significantly decreased at the beginning of the day in the HL-acclimated population relative to the ML control, and the protein was significantly less abundant throughout the day in these cells (Figure 4E). This suggests that intracellular CO_2_ concentrations are likely depleted under diurnal HL, as reported previously for *Chlamydomonas* cells transferred from LL to HL^16^. Low CO_2_ concentration activates the expression of C_i_ transporters, carbonic anhydrases (CAHs), and other genes via the transcription factor CCM1^65,66^. We found that the abundances of LCIB1, LCIC1, and the chloroplast-localized CAH3 and CAH6 were not significantly impacted by diurnal photoacclimation (Figure 4F). These genes are known to be induced under air levels of CO_2_ (∼0.04%), and in our growth conditions (bubbled with ambient air), increased light intensity did not boost their induction further. On the other hand, the chloroplast envelope proteins CCP1 and CCP2; the LCIB1 homologs LCID1 and LCIE1; HLA3, NAR1B, and LCI1, which are known to be induced under very low CO_2_ (< 0.04%); and the mitochondria-localized CAH4 and CAH5 were significantly more abundant in the HL-acclimated population at several timepoints, including in the dark. Thus, it appears that *Chlamydomonas* cells acclimated to diurnal HL are CO_2_ limited and activate their carbon concentrating mechanisms, as reported previously^16^, and we find that this response persists in the nighttime.

### Diurnal photoacclimation leads to differences in thylakoid membranes that persist in the dark phase

Next, we turned our attention to the morphology and composition of the thylakoid membranes, which house the photosynthetic apparatus. In *Chlamydomonas*, thylakoid membranes are organized into appressed stacks of thylakoid membranes and non-appressed, stroma-exposed membranes. The degree of stacking influences the distribution of photosynthetic proteins and the PSII repair machinery, some of which is sterically excluded from the appressed membrane regions where PSII holocomplexes are localized. Therefore, unstacking of thylakoid membranes under HL conditions is important for the degradation of photodamaged D1 subunits and the reassembly of functional PSII holocomplexes^22,67^. In *Chlamydomonas*, the number of membrane layers in appressed regions is reduced in HL^37^, and unstacking occurs within 15 min of increased irradiance^68^. However, it is unknown how thylakoid structure changes over the diurnal cycle and how acclimation to diurnal LL or HL influences stacking in *Chlamydomonas*.

Using transmission electron microscopy (TEM), we found that thylakoid stacks appeared thin and loosely stacked in HL cells across the diurnal cycle (Figures 5A and S5A). Quantitative image analysis revealed that the number of thylakoid layers decreased in the daytime for all three populations but was indeed significantly lower in HL cells than in LL cells at all timepoints, even during the night (Figure 5B, *p* < 0.05). The space between thylakoid layers, measured as the stacking repeat distance, was significantly higher in HL cells than in LL cells throughout the day (*p* < 0.05), and, surprisingly, was highest at the end of the night in the HL cells (Figure 5C, *p-adj.* < 0.05). Thus, acclimation to different intensities of diurnal light resulted in changes to thylakoid structure which persist through the night, suggesting memory or anticipation of the daylight environment.

**Figure 5:**
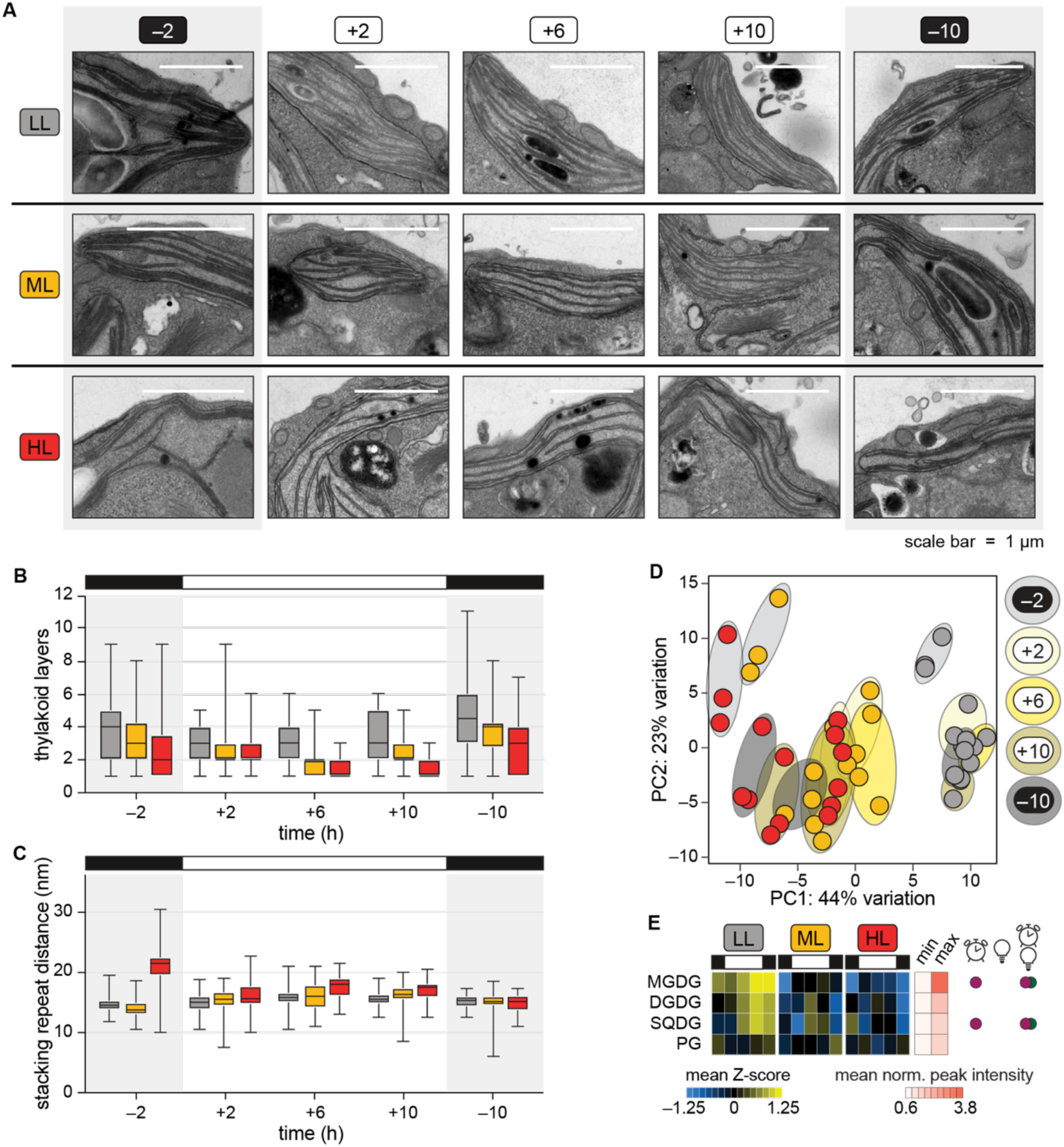
Diurnal photoacclimation leads to differences in thylakoid membranes that persist in the dark phase. (A) Representative electron micrographs of thylakoid membranes. (B) Number of thylakoid membrane layers. (C) Stacking repeat distance of appressed thylakoid membranes. (D) PCA of lipids detected by LC-ESI-MS/MS in positive ionization mode (MGDG, DGDG, SQDG, DG, DGTSA, and TG species); ellipses designate time of day. (E) Average changes in the major classes of chloroplast lipid. Circles at right indicate a significant effect of time, light intensity, or the interaction between the two by two-way mixed ANOVA (*p* < 0.05). See also Figure S5, and Tables S1 and S7.

Our TEM data suggested that total thylakoid membrane area was increased in LL cells and decreased in HL cells. To test this hypothesis and determine whether thylakoid lipid composition is influenced by diurnal light intensity, we monitored the abundance of major chloroplast lipids in the photoacclimated populations using LC-ESI-MS/MS lipidomics (Tables S1 and S7). PCA demonstrated that light intensity had a major influence on the lipidome (Figures 5D and S5C). We found that LL cells had increased levels of thylakoid lipids over the diurnal cycle relative to HL cells, especially monogalactosyldiacylglycerol (MGDG) and sulfoquinovosyldiacylglycerol (SQDG) (Figures 5E, S5B, and S5D). As MGDG and digalactosyldiacylglycerol (DGDG) constitute ∼80% of total thylakoid lipids^69^, our lipidomics data are consistent with the total thylakoid membrane area being increased under diurnal LL. In summary, we found that acclimation to different diurnal light intensities has a major impact on thylakoid architecture and lipid content, causing changes that persist across all stages of the cell cycle.

### Photosystems and their associated LHCs are co-expressed and are decreased across the diurnal cycle in HL-acclimated cells

While PSII is localized to appressed membrane regions, PSI and the chloroplast ATP synthase (CF_o_F_1_) are segregated to non-appressed membranes and the cytochrome (Cyt) *b*_6_*f* complex is distributed throughout the thylakoid membrane system^70,71^ (Figure 6A). We wondered whether the observed light intensity-dependent changes in thylakoid membrane area and stacking had an impact on the accumulation of these complexes. We found that mRNA abundances for all four complexes as well as for the mobile electron carriers plastocyanin, ferredoxin, and the ferredoxin-NADP^+^ reductase peaked in the morning and midday, suggesting coordinated biogenesis independent of the target membrane region (Figures 6B and 6C). The abundances of Cyt *b*_6_*f* and CF_o_F_1_ subunits peaked shortly after their cognate transcripts at the +6 timepoint. Cyt *b*_6_*f* is thought to be a limiting factor in photosynthetic electron transfer, and its increased abundance could contribute to the peak in O_2_ evolution that we observed during the day (Figure 4B). In contrast, the abundances of photosystem subunits were less dynamic overtime. Most PSII and PSI proteins were significantly decreased in HL-acclimated cells throughout the diurnal cycle, including during the dark phase, regardless of their distinct distribution within the thylakoid membrane system (Figure S6A). Plastocyanin and PETO1 were also significantly less abundant in HL-acclimated cells at all timepoints, whereas other subunits of Cyt *b*_6_*f* and CF_o_F_1_ only showed transient significant decreases upon acclimation to diurnal HL (Figure S6B).

**Figure 6:**
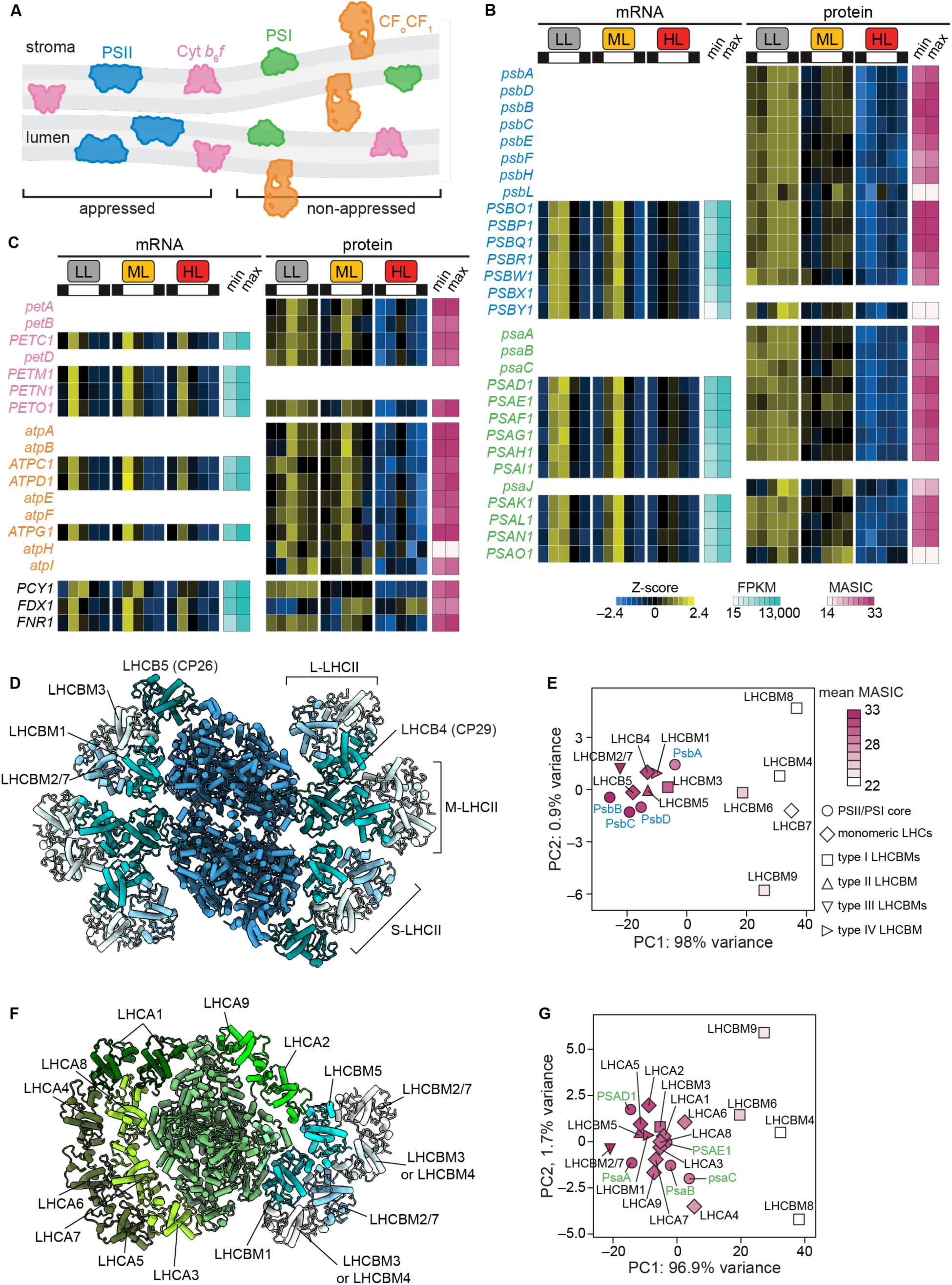
Photosystems and their associated LHCs are co-expressed and are decreased across the diurnal cycle in HL-acclimated cells. (A) Schematic illustrating segregation of PSII to appressed thylakoid membrane regions and of PSI and CF_o_CF_1_ to non-appressed regions. (B) Changes in photosystem mRNAs and proteins. Gene names are colored by complex as in (A). (C) Changes in mRNAs and proteins for other components of the photosynthetic electron transfer chain. Gene names are colored by complex as in (A) or are black for mobile carriers. (D) Structure of the PSII-LHCII supercomplex isolated from *Chlamydomonas* (PDB ID 6KAD). (E) PCA of LHCIIs and PSII core proteins across our proteomics dataset. (F) Structure of the PSI-LHCI-LHCII supercomplex isolated from *Chlamydomonas* (PDB ID 7D0J). (G) PCA of LHCBMs, LHCAs, and PSI core proteins across our proteomics dataset. See also Figure S6.

The *Chlamydomonas* photosystems are surrounded by Chl-binding LHC antenna proteins, which absorb light energy and direct it to the reaction centers. The monomeric LHCII antenna proteins LHCB4 (CP29) and LHCB5 (CP26) bind directly to the PSII core (C) at a 1:1 stoichiometry (Figure 6D). In addition, heterotrimeric LHCIIs composed of the major LHCII proteins, LHCBMs, associate with various affinities to PSII. S-LHCII (strongly associated trimers) attach to LHCB5 and CP43, while the attachment of M-LHCII (moderately associated trimers) and L-LHCII (loosely bound trimers) is mediated by LHCB4^72,73^. Structural analysis of C_2_S_2_M_2_L_2_ PSII supercomplexes isolated from *Chlamydomonas* grown mixotrophically under LL (20–50 µmol photons m^−2^ s^−1^) has revealed that the S-LHCII trimers are primarily composed of LHCBM1, LHCBM2/7 (which are identical proteins encoded by two independent genes), and LHCBM3^72,73^. The composition of the M- and L-LHCII trimers remains to be determined.

We found that the pattern of expression for LHCB4, LHCB5, and LHCBM1–8 mirrored that of the PSII subunits in our experiments: mRNAs accumulated midday, and proteins were decreased in the HL-acclimated cells across the diurnal cycle (Figure S6C). However, some LHCII proteins were much more abundant than others. PCA of these proteins across all samples showed that the monomeric LHCB4 and LHCB5 were co-expressed with the PSII core proteins, as were LHCBM1, LHCBM2/7, LHCBM3, and LHCBM5 (Figure 6E). Each of these LHCBMs has been identified in trimers associated with either PSII or PSI (Figures 6D and 6F)^72,74–79^. PCA also groups these LHCBM proteins with PSI subunits and the nine LHCA antenna proteins, consistent with their recruitment to PSI in State 2 (Figure 6G). Upon phosphorylation by STT7 kinase, LHCBM1 and LHCBM5 engage in a specific interaction with PSI and mediate the binding of LHCII trimers to the PSI core^79^. The recruitment of LHCBM2/7 and LHCBM3 to the PSI-LHCI-LHCII supercomplex may be a function of their high abundance in the thylakoid membrane. The other LHCII proteins (LHCBM4, LHCBM6, LHCBM8, and LHCBM9), which are not known to be present in trimers, were much less abundant on average and were separated from both PSII and PSI core proteins by PCA. Thus, although our proteomics data cannot be used to determine protein stoichiometries, our protein co-expression analysis is consistent with the view that LHCBM1, LHCBM2/7, LHCBM3, and LHCBM5 are the primary antenna proteins present in the LHCII trimers.

### Light stress is dynamic over the daytime and may depend on cell cycle stage

To determine how efficiently absorbed light energy is used for photochemistry, we measured the maximum photochemical efficiency of PSII, or F_v_/F_m_. All three populations exhibited high F_v_/F_m_ (∼0.7) during the night, although it was slightly (but significantly) lower for the HL population (Figure 7A, *p* < 0.05). F_v_/F_m_ of the HL population decreased significantly at the beginning of the day and then improved at the +10 timepoint (*p-adj.* < 0.05). This recovery suggests that the HL cells increase the rate of PSII repair and/or alter their photoprotective strategy to reduce the amount of PSII damage in the latter half of the day.

**Figure 7:**
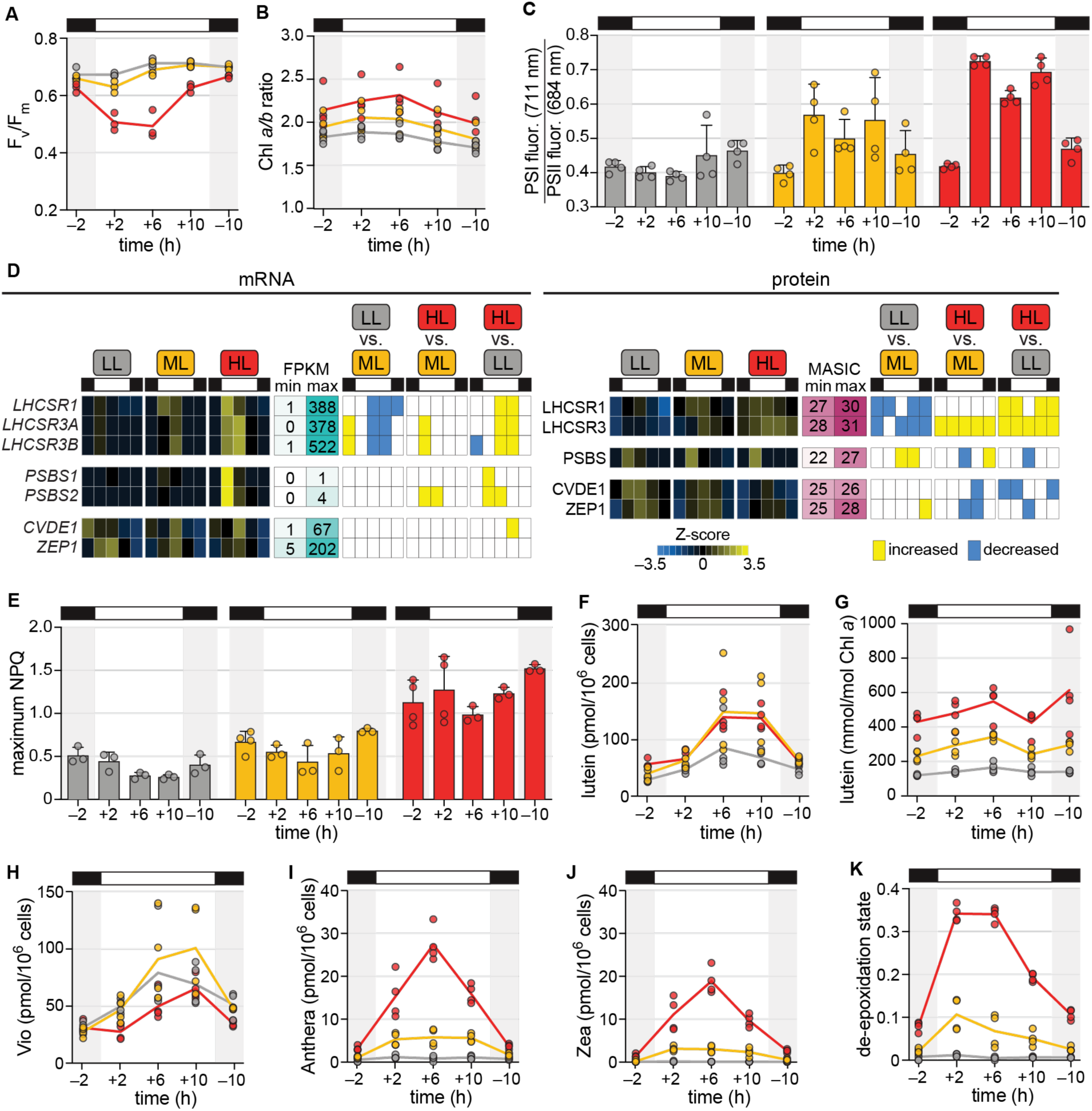
Light stress is dynamic over the daytime. (A) Maximum photochemical efficiency of PSII (F_v_/F_m_). (B) Chlorophyll *a*/*b* ratio. (C) Approximate PSI fluorescence (711 nm) relative to PSII fluorescence (684 nm) measured at 77 K. (D) Changes in mRNA and protein abundance for qE- and qZ-related genes. (E) Maximum NPQ measured from 50–1500 µmol photons m^−2^ s^−1^ actinic light. (F) Cellular lutein content. (G) Lutein content per Chl *a*. (H) Cellular Vio content. (I) Cellular Anthera content. (J) Cellular Zea content. (K) De-epoxidation state of xanthophylls estimated as (0.5Anthera + Zea) / (Vio + Anthera + Zea). See also Figure S7.

As discussed, one way to reduce photodamage is to decrease the size of the light-harvesting antenna. We monitored changes in the ratio of Chl *a* and *b* (Chl *a*/*b*) as a proxy for antenna size, as Chl *b* is exclusively localized to LHC proteins. As compared to the LL population, the HL population exhibited a significantly higher Chl *a*/*b* ratio until the –10 timepoint, suggesting reduced antenna size (Figure 7B, *p* < 0.05). While the Chl *a*/*b* ratio appeared to decrease somewhat from the +6 to the +10 timepoint, this difference was not significant (*p-adj.* > 0.05). On the other hand, we noted that the abundance of LHCBM9 increased substantially at the +10 timepoint in the HL and ML populations, distinguishing it from the other LHC proteins (Figure S6C). LHCBM9 is known to accumulate upon prolonged stress conditions and replace other LHCBM proteins in the antenna, thereby enhancing energy dissipation and stabilizing PSII^80,81^. We hypothesize that increased LHCBM9 at the end of the day could contribute to the recovery in PSII efficiency observed in the HL-acclimated cells.

Phosphorylation by STT7 kinase not only mediates the association of LHCII trimers with PSI during qT, it is also thought to increase electrostatic repulsion between adjacent thylakoid membranes^82–86^, which could contribute to changes in thylakoid structure in HL. To assess the contribution of qT to photosynthetic efficiency and thylakoid morphology, we measured Chl fluorescence emission at 77 K (Figures 7C and S7A). There was almost no change in the spectrum of LL cells over time, indicating that the cells remain in State 1 (more excitation of PSII than PSI). For the ML and HL populations, the PSI fluorescence (∼711 nm) increased relative to PSII fluorescence (∼684 nm) during the day. These changes can indicate not only an increase in the excitation energy transferred to PSI (transition to State 2) but also a decreased amount of PSII fluorescence due to photoinhibition. We also found that the relative PSI fluorescence was significantly decreased at the +6 timepoint compared to the beginning and end of the day for the HL population (*p-adj.* < 0.05). The emission spectra for both the ML and HL cultures returned to the LL-state (State 1) during the night, demonstrating that the observed energy redistribution is completely reversible. Importantly, thylakoid membranes remain less stacked in HL cells relative to LL cells during the night even in the absence of qT (Figure 5). This suggests that LHCII phosphorylation is not necessary for maintaining low thylakoid stacking, as deduced previously from studies of the *Chlamydomonas stt7* kinase mutant^68^. Thylakoid stacking is also mediated by the abundance of LHCII regardless of phosphorylation state^87–91^, so HL-acclimated cells may exhibit lower stacking than LL-acclimated cells in the night by maintaining a smaller population of LHCII proteins in their thylakoid membranes. qT during the day may contribute to the further reduction of stacking during the light phase.

Next, we determined the expression patterns of genes involved in qE. We found that while *LHCSRs* were expressed at all light intensities, mRNA and protein abundance increased with increasing light intensity as expected (Figures 7D). LHCSR1 and LHCSR3 proteins peaked early and midday, respectively, and they remained abundant at the other timepoints examined, as observed previously^41,42^. On the other hand, PSBS only accumulated transiently, and it reached a similar maximum abundance regardless of light intensity. Although PSBS is an essential protein for activating the qE mechanism in land plants^92^, the function of *Chlamydomonas* PSBS orthologs is unclear. We found that NPQ capacity was highest for the population acclimated to diurnal HL, and, in contrast to qT, it was maintained in the dark phase, as previously reported (Figure 7E)^41,42^. Since PSBS levels did not correlate with NPQ capacity, our results support the hypothesis that PSBS is not directly involved in NPQ in *Chlamydomonas*.

It has also been shown that the energy-dissipative carotenoid Zea is not necessary for the qE component of NPQ in *Chlamydomonas*^17^. Lutein, on the other hand, can form a radical cation in the LHCSR3 protein, which could cause qE in the alga^93–95^. We monitored these and other pigments by HPLC (Figures 7F–7J and S7B–S7J). Lutein scaled with cell size over the diurnal cycle and was comparable between the ML and HL populations, as was the case for Vio, ⍺-carotene, and β-carotene (Figures 7F, 7H, S7B, and S7C). Yet, when quantified relative to Chl *a*, lutein abundance was correlated with NPQ capacity, remaining high across the diurnal cycle in the HL-acclimated population (Figure 7G). Antheraxanthin (Anthera) and Zea were significantly higher in the HL cells at all timepoints (Figures 7I, 7J, S7I, and S7J, *p* < 0.05), and unlike lutein, Anthera and Zea peaked midday and then decreased again at the +10 timepoint. As a result, the de-epoxidation state of the HL cells was highest early- and midday, fell substantially at the +10 timepoint, and then remained relatively low during the night (Figure 7K, *p-adj.* < 0.05). This suggests that CVDE1 activity is decreased in the latter half of the day, and since the protein’s activity is responsive to ΔpH, it may indicate that the pH gradient has relaxed at the +10 timepoint. The maintenance of lutein but not high de-epoxidation state over the diurnal cycle provides further evidence that lutein, rather than Zea, is primarily responsible for the qE component of NPQ in *Chlamydomonas*.

Taken together, our results demonstrate dynamic regulation of qT, qE, and qZ to orchestrate a photoprotective quenching strategy tailored not only to the intensity of daylight typically experienced by the cells but also to the time of day.

## DISCUSSION

We have presented an integrative systems analysis of the model unicellular alga *Chlamydomonas* acclimated to challenging light intensities over the diurnal cycle. By monitoring synchronous populations under conditions that simulate our natural environment, we uncovered distinct, physiologically relevant light-responsive changes in gene expression and physiology in both the light and dark phases with high signal-to-noise. We find that the *Chlamydomonas* diurnal program is resilient to both limiting and excess light (Figure 2). Yet, daylight intensity alters the expression of hundreds of genes (Figure 3) and many aspects of the alga’s physiology and metabolism, even during the night.

*Chlamydomonas’* transcriptome and proteome are dynamic over the diurnal cycle, as metabolism and cell cycle progression are coordinated with time of day. Here, we found that the F_v_/F_m_ and de-epoxidation state also change between the beginning, middle, and end of the day even when light intensity remains constant (Figure 7). The F_v_/F_m_ of HL-acclimated cells recovers almost completely at the +10 timepoint, concomitantly with a precipitous drop in the de-epoxidation state. These changes indicate that the HL cells partially overcome light stress once they reach S/M phase of the cell cycle. Studies on marine photosynthetic microorganisms have observed similar recoveries in the afternoon, perhaps speaking to this phenomenon as a common feature of the daily exposure to light stress^96^. Future work should interrogate whether PSII repair activity is enhanced in the latter half of the day, whether LHCBM9 accumulation at this time is required for the observed recovery (Figure S6C), or whether entry into S/M phase itself provides some relief from excess light.

Despite these dynamics in gene expression and photodamage, we found that most photoprotection strategies are constitutive in cells acclimated to diurnal HL. HL-acclimated cells sustained loosely stacked thylakoid membranes and reduced thylakoid lipid content (Figure 5), their photosystems were less abundant (Figures 6 and S6), their Chl content and antenna size were decreased (Figures 1D and 7B), their LHCSR3 abundance and NPQ capacity were increased (Figures 7D and 7E), and their lutein content was higher on a per-Chl basis (Figure 7G) at all timepoints, even after 10 h of dark. These persistent phenotypes suggest a physiological memory of the previous day and may allow the alga to prepare for tomorrow. They also raise questions about how acclimation to one diurnal light intensity impacts fitness and how long it takes for the acclimatory phenotypes to manifest.

Integration of our system-wide measurements on cells acclimated to diurnal HL has contributed to our understanding of the mechanisms of photoprotection in *Chlamydomonas*. For example, our findings suggest that LHCII abundance as well as LHCII phosphorylation both contribute to changes in thylakoid architecture (Figures 5, S6C, and 7C). In addition, our data provide further evidence that PSBS is not directly involved in NPQ in *Chlamydomonas*, and that lutein allows for the constitutive LHCSR-dependent NPQ observed, rather than Zea (Figure 7). To further elucidate light stress responses in conditions that mimic the natural environment, we include a core list of HL-responsive transcripts and proteins at each of the five timepoints examined, comprising 1665 genes, over half of which are not associated with functional annotations (Table S6 and Figure 3C).

In addition to light responses, our transcriptomic and proteomic data also offer insights into macromolecular homeostasis in the eukaryotic cell. For example, they reveal that rhythmically expressed transcripts peak in abundance prior to their cognate proteins, reflecting the central dogma (Figure 2F). Moreover, transient changes in transcript abundances in response to diurnal photoacclimation often generated sustained changes in protein abundances that persisted throughout the day (e.g., Figures 4F, S6, and 7D). Only ∼11% of light-responsive gene expression changes at a given time occurred at both the mRNA and protein levels (Figure S3). While some of these dissimilarities can be attributed to methodological differences in capturing the transcriptome and the proteome, they are also biological^52–55^. Modest correlation between the two is routinely reported^97–104^ as transcripts and proteins are fundamentally different in their form, function, and regulation. Upon a stimulus, mRNA abundance typically changes by orders of magnitude to influence ribosomal occupancy, a phenomenon referred to as superinduction^105,106^. Massive changes in transcript abundance are mediated not only through transcriptional activation and repression but also post-transcriptionally through changes in mRNA half-life, which has been shown to decrease up to 5-fold in response to light^48–51^. The dynamic range of protein abundances is typically much smaller than that of their cognate transcripts, as protein abundance is tightly constrained to maintain proteostasis. Since cell volume roughly doubles over the diurnal cycle in our experiments, the absolute abundance of a protein would also only need to double to maintain its cellular concentration. Proteins are more often regulated by posttranslational modifications: phosphorylation^107–109^, reduction/oxidation of cysteines^110–112^, acetylation of lysines^113^, and protonation of side chains^94,114,115^ induce rapid, reversible changes in conformation, enzyme activity, protein-protein interactions, and localization in response to changing fluence and other stimuli. Our genome-wide data and paired physiological measurements hint at widespread posttranscriptional regulation specific to different times of day and in response to different light intensities waiting to be uncovered.

## METHODS

### Strains and culture conditions

*Chlamydomonas reinhardtii* strain CC-5390 [CC-4351 (*cw*15–325 *mt*+) rescued with the pCB412 cosmid carrying the *ARG7* gene] was used for all experiments. Cells were precultivated in 125 mL Erlenmeyer flasks containing 50 mL Tris-acetate-phosphate (TAP) medium with a modified trace element solution^116^ in an Innova incubator (New Brunswick Scientific, NJ, USA) with constant agitation at 140 rpm, 24 °C, and continuous white light (50–60 μmol photons m^−2^ s^−1^). Cultures were synchronized and grown in presterilized turbidostat flat-panel FMT 150 Photobioreactors (Photon Systems Instruments, Drásov, Czechia) containing high-salt (HS) medium supplemented with a modified trace element solution^116,117^ as described previously^41^. Briefly, photobioreactors were inoculated with cells precultivated in TAP to a starting optical density (OD_680_) of ∼0.05. Cultures were aerated with filter-sterilized ambient air provided by an aquarium pump, which also mixed cultures with airlift. Cells were grown under 12-h-light/12-h-dark cycles, where temperature was set to 28 °C during the light phase and 18 °C during the dark phase, and illumination was provided by a panel of red and blue LEDs (80% blue, 20% red, LED Light Source SL 3500; Photon Systems Instruments, Drásov, Czechia). Light intensity was set to 50 μmol photons m^−2^ s^−1^ for LL, 200 μmol photons m^−2^ s^−1^ for ML, and 1000 μmol photons m^−2^ s^−1^ for HL. Cells were acclimated to the respective diurnal light condition by growing in turbidostat mode for a minimum of 2 weeks before performing the experiments. As the LL condition limited growth, cells were first grown for 1 week at ML, and then the light was changed to LL for at least 1 week before performing the experiments. At the start of each experiment, OD_680_ was set to 0.4 at the beginning of the dark phase, and the flow of media was stopped in order to monitor changes in culture density. Experimental replicates refer to independent experiments performed with independent cultures in different weeks.

### Live-cell imaging by Airyscan microscopy

10 mL of culture was collected by centrifugation at 1800 x*g* for 1 min, and cells were resuspended in 0.7% low-melting-point agarose in HS medium and mounted between slide and coverslip. Cells were observed using a Zeiss LSM 880 or a Zeiss LSM 980 microscope equipped with the Airyscan detector with a Zeiss Plan-Apochromat 63×/1.4 NA DIC M27 Oil objective (Zeiss, Oberkochen, Germany). Chlorophyll (Chl) was excited with a 633 nm laser, and fluorescence was acquired through a 645 nm longpass filter. At least 12 representative cells were imaged per condition, and one representative image was chosen for display. Image acquisition and analysis were performed using ZEN software (Zeiss, Oberkochen, Germany) and FIJI image analysis software^118,119^.

### Cell density, number, and size

Cell density was continuously monitored by the bioreactors as OD_680_, and the mean OD_680_ and standard deviation across 9 experimental replicates (n = 9) is displayed. Cell number was determined for all experimental replicates with a Z2 Coulter Particle Count and Size Analyzer (Beckman Coulter, CA, USA) set to count particles between 4–14 µm in diameter. Cell diameter was determined for 3 experimental replicates (n = 3). Images of cultures immobilized with iodine were taken <15 min after each timepoint at 100X total magnification using ZEN software and diameters were estimated using FIJI image analysis software. In FIJI, images were made binary, holes were filled, and the “watershed” command was used to distinguish actively dividing daughter cells and count them as individual cells. The conversion factor of 0.44 µm/pixel (corresponding to the magnification of the image) was used to determine the diameter of particles ranging from 15– 3000 µm^2^ in area with a circularity factor from 0.5–1. The diameter of 100 cells from each of the 3 experimental replicates (300 cells total) is plotted for each condition.

### Pigment content

Pigment content was measured by HPLC. 10 mL of culture from each of 4 experimental replicates (n = 4) was collected by centrifugation at 2400 x*g* for 3 min at 4 °C, and the pellets were flash-frozen in liquid N_2_ and stored at –80 °C until further processing. After thawing, 100 µL of 100% acetone was used to extract pigments from each sample by vortex for 10 min. The sample was centrifuged at 21,000 x*g* for 5 min at 4 °C, and the supernatant was transferred to a collection tube. Another 100 µL of 100% acetone was added to the pellet to extract pigment completely by vortexing for 10 min. The pellet was removed by centrifugation at 21,000 x*g* for 5 min at 4 °C, and the supernatant was added to the same collection tube as before. The extracted pigments were passed through a 0.22 µm nylon filter and transferred to an HPLC sample vial. HPLC was performed using a Spherisorb ODS1 C_18_ column (4.6 × 250 mm, Waters Corporation, MA, USA) and pigment analysis was performed as previously described^120^. Briefly, pigments were eluted with a linear gradient from 100% (v/v) solvent A [84% (v/v) acetonitrile, 2% (v/v) methanol, 14% (v/v) 0.1 M Tris-HCl (pH 8.0)] to 100% (v/v) solvent B [68% (v/v) methanol, 32% (v/v) ethyl acetate] for 15 min, followed by 3 min of solvent B. The solvent flow rate was 1.0 mL/min. Pigments were detected at 445 nm with a reference at 550 nm by a diode array detector. Pigment content was normalized to either cell number or Chl *a*.

### Total organic carbon measurement

Total nonpurgeable organic carbon (NPOC) of cells was determined for 3 experimental replicates (n = 3) using a TOC-L Shimadzu Total Organic Carbon Analyzer (Shimadzu, Kyoto, Japan). 13 mL of culture was collected by centrifugation at 3200 x*g* and 4 °C for 2 min, and the cell pellets were washed once in 10 mM Na-phosphate buffer (pH 7.0). Cell pellets were stored at –20 °C until analysis (< 1 month). To digest cells, pellets were resuspended in 3 M HCl to achieve roughly 3.33 × 10^7^ cells/mL acid and then incubated at 65 °C for 48 h with constant agitation. Cell digests were diluted 111-fold with MilliQ H_2_O for a final HCl concentration of 27 mM. All samples were sparged to remove inorganic carbon; some organic carbon may be purged from the sample by this method, so we report “nonpurgeable” organic carbon.

### RNA extraction

15 mL of culture from each of 3 experimental replicates (n = 3) was collected by centrifugation at 2500 x*g* and 4 °C for 5 min. Cell pellets were resuspended in 1mL lysis buffer containing 50 mM Tris-HCl pH 7.5, 150 mM NaCl, 15 mM EDTA pH 8, 2% (w/v) SDS, and 40 µg/mL Proteinase K (from *Tritirachium album*, Fisher Scientific, NH, USA) and mixed by gentle pipetting and inversion for 30 s. Cellular starch was removed by centrifugation at 600 x*g* for 3 min, and the lysate was transferred to a fresh 15 mL tube and flash-frozen in liquid N_2_ before storing at –80 °C. RNA from 4–5 frozen samples was extracted in parallel; samples were kept on ice and solutions were pre-cooled when possible. 10 mL TRIzol (Invitrogen, Thermo Fisher Scientific, MA, USA) was added to each frozen pellet and samples were gently inverted until thawed. Then, nucleic acids were extracted by adding 2 mL chloroform/isoamylalcohol (24:1, v/v) and shaking vigorously for 30 s. The sample was quickly transferred to MaXtract High Density tubes (QIAGEN, MD, USA), phases were separated by centrifugation at 1500 x*g* and 20 °C for 5 min, and the aqueous phase was carefully transferred to a fresh tube and mixed with ∼10 mL cold ethanol (100%). The mixture was transferred onto an RNAeasy Mini column (QIAGEN, MD, USA) using a vacuum manifold, washed according to manufacturer’s instructions but with buffer RWT instead of RW1, DNase treated on the column, washed with buffer RWT and RPE, and eluted into 90 µL RNase-free water. In an additional step, the RNA was further purified by precipitation with 0.3 M sodium acetate (pH 7.0) and 2.5-volumes of ethanol (100%) for 30 min at –20 °C, followed by centrifugation at 16,200 x*g* and 4 °C for 30 min. The resulting pellets were washed with 70% (v/v) ethanol and resuspended in 60 µL RNAse-free water. The concentration of total RNA was estimated using a NanoDrop One (Thermo Fisher Scientific, MA, USA) to be >100 ng/µL. RNA quality was assessed by the ratios of A_260_/A_280_ ∼2.05–2.20 and A_260_/A_230_ >2.00, and by using the Agilent RNA 6000 Pico Kit with a 2100 Bioanalyzer according to the manufacturer’s instructions (Agilent, CA, USA).

### Transcriptomics sequencing and quantitation

Total RNA was subjected to poly(A) selection, RNA-Seq library construction, and sequencing on the Illumina HiSeq 3000 platform by the University of California Los Angeles Technology Center for Genomics and Bioinformatics (UCLA, CA, USA) using standard kits and protocols (Illumina Inc. CA, USA). Sequencing comprised 3 lanes of 50 nt reads, which generated 933 M reads total from 45 libraries (three light intensities x five timepoints x three experimental replicates). The resulting reads were mapped to the *C. reinhardtii* reference genome assembly and annotations v6.1 (Phytozome) with STAR (v2.4.0j) using --runThreadN 4 --alignIntronMax 5000. The resulting sam formatted files were compressed, sorted and indexed with samtools (v1.16.1) using view -b -h, sort, and index, respectively. FPKMs were calculated with cuffdiff (v2.0.2) using --max-bundle-frags 1000000000 --library-type fr-firststrand. FPKM values for all 16,735 nucleus-encoded mRNAs detected are available in Table S2.

### Protein and lipid extraction

35 mL of culture from each of 3 experimental replicates (n = 3) was collected by centrifugation at 2400 x*g* at 4 °C for 3 min, resuspended in 200 µL Na-phosphate buffer (pH 7.0), transferred to 1.5 mL tubes, flash-frozen in liquid N_2_, and stored at –80 °C. Cell pellets were extracted using the Metabolite, Protein, and Lipid extraction (MPLex) method^121,122^. The cell pellets were resuspended in Type 1 water and cold (–20 °C) chloroform/methanol (2:1, v/v) in chloroform-compatible 2 mL Sorenson MµlTI™ SafeSeal™ microcentrifuge tubes (Sorenson BioScience, UT, USA). The resulting mixture had a final ratio of 8 parts chloroform, 4 parts methanol, 3 parts water/sample. Samples were vortexed, sonicated on ice in a bath sonicator, and then incubated in an ice block with shaking for 45 min. Samples were then sonicated with a 3 mm probe (20 kHz fixed ultrasonic frequency, sonicator model FB505, Thermo Fisher Scientific, MA, USA) at 20% of the maximum amplitude for 30 s on ice within a fume hood. The polar and nonpolar phases, as well as the protein interlayer, were separated by centrifugation at 12,000 x*g* for 10 min.

The upper polar phase was removed. The protein interlayer was washed with cold 100% methanol, resuspended by vortexing, and the protein was collected by centrifugation at 12,000 x*g* for 5 min. Protein pellets were lightly dried under an N_2_ stream and stored at –80 °C until analysis. The lower nonpolar phase was collected for lipidomics analysis: the solvent was removed by drying in a speed vac, 500 µL of cold chloroform/methanol (2:1, v/v) was added, and the samples were stored at –20 °C until analysis.

### Protein digestion

8 M urea was added to each protein pellet, and a bicinchoninic acid (BCA) assay was used to determine the protein concentration (Thermo Fisher Scientific, MA, USA). Then, 10 mM dithiothreitol (DTT) was incorporated into the samples by incubating at 60 °C for 30 min with continuous agitation at 800 rpm. The samples were then diluted 8-fold before digestion in a solution containing 100 mM triethylamonium bicarbonate (TEAB), 1 mM CaCl_2_, and sequencing-grade modified porcine trypsin (Promega, WI, USA) at a trypsin-to-protein ratio of 1:50 (w/w) for 3 h at 37 °C.

Following digestion, the samples were desalted using a 4-probe positive pressure Gilson GX-274 ASPEC™ system (Gilson Inc., WI, USA) with Discovery C18 100 mg/1 mL solid phase extraction columns (Supelco, MO, USA). Initial conditioning was performed with 3 mL of methanol followed by 2 mL of 0.1% trifluoroacetic acid (TFA) in H_2_O, the samples were loaded onto each column, washed with 4 mL of a solution of 95:5 H_2_O:acetonitrile and 0.1% TFA, and eluted with 1 mL of 80:20 acetonitrile:H_2_O containing 0.1% TFA. The eluted samples were concentrated to approximately 100 µL using a speed vac, and another BCA assay was performed to determine the peptide concentration. 2 µg from each sample was combined to form a global pool, while 30 µg from each sample was aliquoted into new tubes for isobaric multiplexing. These samples were completely dried in a speed vac.

### Tandem mass tag (TMT) peptide labeling

Samples were pooled randomly into 3 groups called “plexes.” Each plex also contained a global pool for normalization across sets. Dried samples were diluted in 40 µL of 500 mM HEPES (pH 8.5) and labeled using an amine-reactive Tandem Mass Tag Isobaric Mass Tagging Kit (Thermo Fisher Scientific, MA, USA) according to the manufacturer’s instructions. 250 μL of anhydrous acetonitrile was added to each 5 mg reagent, which was dissolved over 5 min with occasional vortexing. Reagents (10 µL) were then added to each sample and incubated for 1 h at ambient temperature with shaking at 400 rpm. Each sample was then diluted with 30 µl 20% acetonitrile (v/v). A portion from each sample was collected as a premix, which was run on a mass spectrometer to confirm complete labeling. The samples were frozen at –80 °C until further analysis. Frozen samples were thawed, and the reaction was quenched by adding 8 μL of 5% hydroxylamine (v/v) and incubating for 15 min at ambient temperature with shaking at 400 rpm. The samples within each set were then combined and completely dried in a speed vac. Samples were then cleaned using Discovery C18 50 mg/1 mL solid phase extraction tubes as described above and once again assayed by BCA to determine the final peptide concentration. Samples were diluted in 2% ACN, 0.1% formic acid to a peptide concentration of 0.25 μg/μL for microfractionation.

Peptide mixtures were separated by high resolution reversed phase UPLC using a nanoACQUITY UPLC® system (Waters Corporation, MA, USA) equipped with an autosampler. Capillary columns (200 µm i.d. × 65 cm long) were packed with 300 Å 3.0 µm p.s. Jupiter C18 bonded particles (Phenomenex, CA, USA). Separations were performed at a flow rate of 2.2 µL/min on binary pump systems using 10 mM ammonium formate (pH 7.5) as mobile phase A and 100% acetonitrile as mobile phase B. 48 µL of TMT labeled peptide mixtures (0.25 µg/µL) were loaded onto the column and separated using a binary gradient of 1% B for 35 min, 1–10% B in 2 min, 10–15% B in 5 min, and 15–25% B in 35 min, 25–32% B in 25 min, 32–40% B in 13 min, 40–90% B in 43 min, held at 90% B for 2 min (column washed and equilibrated from 90–50% B in 2 min, 50–95% B in 2 min, held at 95% B for 2 min and 95–1% in 4 min). The capillary column eluent was automatically deposited every minute into 12 × 32 mm polypropylene vials (Waters Corporation, MA, USA) starting at 60 min and ending at 170 min over the course of the 180 min LC run. Fractions were concatenated by collecting fractions from vial 1 to vial 24 and then returning to vial 1 back to vial 24 (etc.). Prior to peptide fraction collection, 100 µL of water was added to each vial to aid the addition of small droplets into the vial. Each vial was completely dried in a vacuum concentrator (Labconco, MO, USA) and reconstituted in 20 µL 2% acetonitrile, 0.1% formic acid and stored at –20 °C until LC-MS/MS analysis.

### TMT-labeled peptide mass spectrometry

A Thermo Dionex Ultimate 3000 UPLC system (Thermo Fisher Scientific, MA, USA) was configured to load a 5 µL injection directly onto the column at a flow rate of 200 nL/min, allowing 40 min to load the sample onto the column before the elution gradient was started. The analytical column was made using an integrated emitter capillary (75 µm i.d. x 25 cm long), packed in-house using BEH C18 media (Waters Corporation, MA, USA) in 1.7 µm particle size. Columns were heated to 45 °C using the MonoSLEEVE controller and a 15 cm heater (Analytical Sales and Services Inc., NJ, USA). Mobile phases consisted of (A) 0.1% formic acid in H_2_O and (B) 0.1% formic acid in acetonitrile with the following gradient profile (min, %B): 0, 1; 40, 1; 50, 8; 145, 25; 155, 35; 160, 75; 165, 5; 170, 95; 175, 1.

MS analysis was performed using a Q Exactive HF-X mass spectrometer (Thermo Fisher Scientific, MA, USA) outfitted with a Nanospray Flex™ Ion Source (Thermo Fisher Scientific, MA, USA) ionization interface. The ion transfer tube temperature and spray voltage were 300 °C and 2.2 kV, respectively. Data were collected for 120 min following a 60 min delay from sample injection. FT-MS spectra were acquired from 300–1800 m/z at a resolution of 60 K (automatic gain control (AGC) target 3 x10^6^) while the top 12 FT-HCD-MS/MS spectra were acquired in data-dependent mode with an isolation window of 0.7 m/z and a resolution of 45 K (AGC target 1 x10^5^) using a normalized collision energy of 30% and an exclusion time of 45 s.

### TMT proteomics data analysis

LC-MS/MS datafiles were converted to mzML format using MSConvert, part of the ProteoWizard^123^ suite of software tools. MZRefinery^124^ was employed to re-calibrate the MS level search mass values. Adjusted spectra were then searched using MS-GF+^125^ with partial tryptic cleavage rules, +/– 20 ppm parent mass tolerance, static TMT 16-plex modification (+229.1629 Da) of peptide N-termini and lysine residues, dynamic xxidation (+15.9949 Da) of methionine residues, and instrument set to Q-Exactive derived data. A target/decoy approach was employed^126^ against 32,670 protein sequences from the *Chlamydomonas reinhardtii* v6.1 genome annotation^127^ coupled with 16 commonly observed contaminants (trypsin, keratin, etc.). Peptide-to-spectrum matching (PSM) identifications were filtered to 1% FDR (10,277 decoy from 1,027,720 filter passing PSMs), matched back to the search FASTA using the Protein Coverage Summarizer tool^128^, and peptide uniqueness was assessed. It was determined that using Gene IDs for peptide grouping was preferred (e.g., Cre01.g000100_4532.1.v6.1 and Cre01.g000100_4532.2.v6.1 group to Cre01.g000100), and parsimony was applied such that only non-unique peptides mapping to more than one uniquely identified Gene IDs (2.1% of the overall data) were excluded from quantitation. The MASIC software tool^129^ was used to extract TMT reporter ion signal abundances within a tolerance of +/– 0.003 Da, as well as an isotopic interference assessment for precursor signals. Signals with more than 20% non-parent isotopic interference (≤ 0.8) were excluded from quantitation. Resulting peptides and their associated abundances (MASIC values) were log_2_-transformed, visualized, and mean central tendency normalized using InfernoRDN^130^. Normalized peptide abundances were de-logged, grouped by parsimony-filtered Gene IDs, and abundances summed (simple protein rollup). Resulting Gene ID abundances were log_2_-transformed and mean central tendency normalized to remove small, experimentally induced variations, resulting in the final MASIC values reported.

While 11,462 proteins were detected across the dataset, only proteins which were detected in at least two of the three experimental replicates at all timepoints were analyzed (10,075 proteins total); MASIC values for these proteins are available in Table S2.

### Enzymatic starch measurement

20 mL of culture from each of 3 experimental replicates (n = 3) was collected in Protein LoBind tubes (Eppendorf, Hamburg, Germany) by centrifugation at 1650 x*g* for 10 min. Cell pellets were flash frozen in liquid N_2_ and then stored at –80 °C until processing (< 2 weeks). Starch was extracted from the cell pellets by ethanolic extraction in three subsequent cycles. Each pellet was resuspended in 250 µL 80% (v/v) ethanol, and the suspensions were transferred to screw-cap tubes with rubber gaskets to minimize evaporation loss. Each tube was incubated at 95 °C for 30 min with constant agitation, allowed to cool to ambient temperature (∼5 min), and the pellet collected by centrifugation at 5000 x*g* for 10 min. The supernatant was carefully removed, and the pellet was subsequently treated in the same way with 150 µL of 80% (v/v) ethanol and then with 250 µL of 50% (v/v) ethanol, respectively. The resulting pellets were hydrolyzed by resuspending in 400 µL of 0.1 M NaOH, shaking vigorously, incubating at 95 °C for 30 min with constant agitation, allowed to cool to ambient temperature, and the pellet collected by centrifugation at 5000 x*g* for 10 min. The supernatant was carefully removed, and the pellet was resuspended in 80 µL of freshly prepared neutralization solution (0.1 M sodium acetate, with pH adjusted to 4.9 with NaOH and 0.5 M HCl) and 320 µL of MilliQ H_2_O was added to prepare the final sample volume. The starch content in each sample was determined using a Starch Assay Kit (Sigma-Adrich, MO, USA) according to the manufacturer’s instructions, but with the sample preparation replaced by the ethanolic extraction and hydrolysis described above, and the assay adjusted to smaller reaction volumes. The starch assay volume was reduced to 350 µL while maintaining reagent ratio and the glucose assay was performed in 96-well plates using 20 µL of the starch assay reaction in a total reaction volume of 200 µL. Cellular glucose levels were determined for each sample in a parallel reaction lacking amyloglucosidase, and the resulting background glucose concentration was subtracted from the final glucose concentration (cellular glucose + glucose derived from starch) to calculate starch content. Starch concentration was normalized to cell number.

### O_2_ consumption and evolution measurements

O_2_ consumption and evolution rates were measured for 3 experimental replicates (n = 3) on a standard Clark-type electrode (Hansatech Oxygraph with a DW-2/2 electrode chamber) and analyzed with Hansatech O2view software (v2.10, Hansatech Instruments Ltd, Norfolk, UK). 2 mL of cells sampled at the designated timepoints were assayed in the presence of 5 mM NaHCO_3_ under constant agitation. Respiration rates were measured as O_2_ consumption over a period of 5 min in the dark. Then, O_2_ evolution was measured during illumination with 50 μmol photons m^−2^ s^−1^ for 5 min, 200 μmol photons m^−2^ s^−1^ for 3 min, and 1000 μmol photons m^−2^ s^−1^ for 3 min. As O_2_ evolution rates were highest during illumination with 1000 μmol photons m^−2^ s^−1^, only those rates were reported. The rate of photosynthetic O_2_ evolution was calculated as the difference between O_2_ evolution and O_2_ consumption in the dark for each sample. O_2_ consumption and evolution rates were normalized to NPOC as a proxy for biomass or to Chl *a*.

### Transmission electron microscopy (TEM)

50 mL of culture was collected by centrifugation at 1800 x*g* for 1 min, cells were resuspended in 2% glutaraldehyde in HS medium, and fixed by incubation at 4 °C with rotatory agitation in the dark for > 10 h. The fixed cells were washed and post-fixed with Na-cacodylate buffer containing 1% (w/v) OsO_4_ and 1.6% (w/v) potassium ferricyanide for 1 h. The fixed cells were washed twice with cacodylate buffer for 10 min. The washed cells were then dehydrated with increasing concentrations of acetone (35–100%). Dehydrated cells were infiltrated and embedded in EPON resin. Sections were cut to approximately 70–100 nm in thickness. The thin sections were collected onto copper formvar slot grids. Sections were post-stained with 2% uranyl acetate for 7 min, followed by lead citrate for 7 min. The sections were dried and examined using a JEOL 1200 EX 100 kV TEM (JEOL, MA, USA) with an Orius CCD camera (Gatan Inc., CA, USA) or a Technai 12 120 kV FEI TEM with a Rio16 CMOS Camera (Gatan Inc, CA, USA). At least 15 representative cells were imaged from each sample, and one representative cell was chosen for display.

### Image analysis of thylakoid membranes

The characteristics of thylakoid membranes [number of thylakoid membrane layers, membrane height, and stacking repeat distance (SRD)] were manually measured for at least 37 representative thylakoid membrane regions per sample (n ≥ 37) (from at least 10 representative cells per sample) using FIJI image analysis software as previously described^131^. The analysis method was modified as follows because *Chlamydomonas* thylakoid membrane architecture has less distinct grana structures than do land plants. Instead of only collecting data on thylakoid membrane stacks with three or more membrane layers, we collected data on all levels of stacking, including single, unstacked thylakoid membranes. Membrane height is defined as the distance between the top and bottom layers of the outer thylakoid membranes of a stack. SRD represents the average thickness of each thylakoid membrane and is calculated by dividing the height of the membrane stack by the number of membrane layers in the stack. SRD was only determined for stacked thylakoid membranes (2 or more thylakoid membrane layers).

### Lipidomics analysis

Untargeted lipid detection, identification, and quantitation were achieved by LC-ESI-MS/MS analysis using a Vanquish Flex UHPLC system (Thermo Fisher Scientific, MA, USA) interfaced with a Velos-ETD Orbitrap mass spectrometer (Thermo Fisher Scientific, MA, USA). Dried lipids extracted by MPLex (described above) were reconstituted in 90% methanol and 10% chloroform (v/v) and injected onto an ACQUITY UPLC CSH column (3.0 mm × 150 mm × 1.7 µm particle size, Waters Corporation, MA, USA). Lipids were separated over a 34 min gradient elution (mobile phase A: acetonitrile:H_2_O (40:60, v/v) containing 10 mM ammonium acetate; mobile phase B: acetonitrile:isopropyl alcohol (10:90, v/v) containing 10 mM ammonium acetate) at a flow rate of 250 µL/min. The full gradient profile was as follows (min, %B): 0, 40; 2, 50; 3, 60; 12, 70; 15, 75; 17, 78; 19, 85; 22, 92; 25, 99; 34, 99; 34.5, 40. The mass spectrometer inlet and HESI source were maintained at 350 °C with a spray voltage of 3.5 kV and sheath, auxiliary, and sweep gas flows of 45, 30, and 2, respectively. Samples were analyzed in both positive and negative ionization using higher-energy collision dissociation (HCD) and collision-induced dissociation (CID) to obtain high coverage of the lipidome. Data were collected using a precursor scan of 200– 2000 m/z at a mass resolution of 60 k, followed by data-dependent MS/MS of the top 4 ions. Normalized collision energies for CID and HCD were 35 and 30, respectively. CID spectra were acquired in the ion trap using an activation Q value of 0.18, while HCD spectra were acquired in the orbitrap at a mass resolution of 7.5 k. LC-MS/MS raw data files were imported into the in-house developed software LIQUID (Lipid Informed Quantitation and Identification)^132^ for semi-automated identification of lipid molecular species. Confident lipid identifications were determined by examining the tandem mass spectra for diagnostic ion fragments along with associated chain fragment information. In addition, the isotopic profile, extracted ion chromatogram (XIC), and mass error of measured precursor ions were examined for lipid species. The identified lipid name, observed m/z, and the retention time from each analysis was used as the target database for feature identification across all LC-MS/MS runs to align and gap-fill the mass spectrometry data. This was achieved by aligning all datasets (grouped by ionization type) and matching unidentified features to their identified counterparts using MZmine 2^133^. Aligned features were manually verified, and peak intensity values were exported for statistical analysis^134^.

268 lipid species were detected: 199 in positive mode (DG, DGDG, DGTSA, MGDG, SQDG, & TG species) and 69 in negative mode (PA, PE, PG, & PI species). Peak intensity was log_2_-transformed and then median-normalized to correct for differences in material loading across samples. Median-normalized peak intensity for all lipid species detected are available as Table S7.

### F_v_/F_m_ measurement

F_v_/F_m_ was determined for each of 3 experimental replicates (n = 3). Roughly 10 mL of culture was collected into 25 mL Erlenmeyer flasks and dark-adapted for 15 min. Then, 350 µL of culture was aliquoted into each of 18 replicate wells of a 96-well plate per sample to measure maximum quantum efficiency of PSII using an IMAG-MAX/L MAXI Imaging PAM system equipped with ImagingWinGigE software (Heinz Walz GmbH, Effeltrich, Germany). F_v_/F_m_ was calculated as F_v_/F_m_ = (F_m_ – F_o_)/F_m_, where F_m_ is the maximum fluorescence measured immediately after a saturating pulse and F_o_ is the initial fluorescence of dark-adapted cells. The saturating pulse was administered at an intensity of 10 units and a pulse width of 0.48 s. Gain and damping of the instrument were set to 18 and 4, respectively, such that the F_t_ of a dark-adapted control culture was between 0.15–0.20. Control cultures were grown in TAP medium under 50–60 µmol photons m^−2^ s^−1^ (mixture of warm and cool white light) and 24 °C with constant agitation at 140 rpm in an Innova incubator, and were then kept at a lower light intensity for the duration of each experiment time course (∼20 µmol photons m^−2^ s^−1^).

### 77 K fluorescence emission spectra

Steady-state Chl fluorescence emission spectra at 77 K were measured for 4 experimental replicates (n = 4) with a FluoroMax-4 spectrophotometer (Horiba Scientific, Kyoto, Japan). 1.5 mL of culture was sampled into glass tubes and immediately frozen by submerging in liquid N_2_. The excitation wavelength was 420 nm with a 2 nm slit size. Fluorescence emission was measured from 650 to 780 nm with a 4 nm slit size. Fluorescence emission at each wavelength was measured for 0.1 s, the spectra were recorded three consecutive times for each sample, and the averaged spectra for each replicate sample were obtained. Fluorescence at each wavelength was normalized relative to the maximum fluorescence (resulting from PSII, ∼684 nm). The mean spectra and standard deviation across the 4 experimental replicates are displayed. The PSI fluorescence relative to the PSII fluorescence is reported as the ratio between the fluorescence at the 711 nm peak and the fluorescence at the 684 nm peak.

### Non-photochemical quenching (NPQ) capacity

NPQ capacity was determined by monitoring Chl fluorescence kinetics for at least 3 experimental replicates (n = 3). 20 mL of culture was collected into 50 mL Erlenmeyer flasks and dark-acclimated for 30 min with constant agitation at 180 rpm to relax the proton gradient across the thylakoid membrane and allow for full conversion of Zea to Vio. Agitation was used to prevent hypoxia, which is known to induce State 2 in *Chlamydomonas*^135^. 8–10 mL of culture was then deposited onto a glass fiber prefilter (Millipore Sigma, MA, USA) using a syringe, the filter was positioned in a leaf clip, and light curves were collected with a Fluorescence Monitoring System FMS 2+ (Hansatech Instruments Ltd, Norfolk, UK) as described previously^136^ with gain set to 50 and modulation beam intensity set to 2. The samples were first illuminated with far-red light to induce State 1: samples collected during the light phase were illuminated with far-red light for 10 min, and samples collected during the dark phase were illuminated for 5 min since 77 K fluorescence emission spectra showed that cells were in State 1 in the night. Then, F_v_/F_m_ was measured using a saturating pulse with an intensity of 45 units and a pulse width of 0.5 s. 5 min treatments of increasing actinic light intensity (50, 200, 500, 1000, and 1500 µmol photons m^−2^ s^−1^) were then administered to determine light-adapted F_m_’ (using the same saturating pulse settings) and calculate energy dissipation as NPQ = (F_m_ – F_m_’)/F_m_’.

### Statistical analysis of physiological data

The various physiological data (optical density, cell density, cell size, pigment content, NPOC content, starch content, O_2_ consumption, O_2_ evolution, F_v_/F_m_, and NPQ capacity) were tested for significant differences across at least 3 experimental replicates (n = 3) using Student’s t-tests as follows. Differences between the three populations (acclimated to LL, ML, and HL) at a given time point were tested for significance using two-tailed Student’s t-tests (*p* < 0.05) in Microsoft Excel. Differences between each of the five timepoints (–2, +2, +6, +10, –10) in a given population (LL, ML, or HL) were tested for significance using pairwise t-tests in the R package *rstatix* (v0.7.2) with a Benjamini Hochberg *p-*value adjustment (*p-adj.* < 0.05). Experimental replicates refer to independent experiments performed in different weeks. The number of experimental replicates for each physiological measurement is specified in the associated section of the Methods.

### Statistical analysis of imaging data

For Airyscan microscopy and TEM, at least 12 representative cells were imaged for each condition, and a representative cell was chosen for display. Differences in thylakoid membrane stacking were tested for significant differences across at least 37 representative thylakoid membrane regions per condition (n ≥ 37) (from at least 15 representative cells per condition) using Student’s t-tests as follows. Differences between the three populations (acclimated to LL, ML, and HL) at a given time point were tested for significance using two-tailed Student’s t-tests (*p* < 0.05) in Microsoft Excel. Differences between each of the five timepoints (–2, +2, +6, +10, –10) in a given population (LL, ML, or HL) were tested for significance using pairwise t-tests in the R package *rstatix* (v0.7.2) with a Benjamini Hochberg *p-*value adjustment (*p-adj.* < 0.05).

### Significant differences in the transcriptome

Differences in mRNA abundance between the three populations (acclimated to LL, ML, and HL) at a given time point were tested for significance with the R package *DESeq2* (v1.44.0). Transcript abundance differences were considered significant if they had a log_2_-transformed fold change > 1 or < –1 and a Benjamini-Hochberg corrected *p*-value < 0.01. The full list of significant changes in mRNA abundance is available in Table S5. Significant differences between the three populations at a given time are represented as yellow or blue tiles in figures.

### Significant differences in the proteome

Differences in protein abundance between the test populations (acclimated to LL or HL) relative to the control population (acclimated to ML) at a given time point were tested for significance as follows. The average MASIC value for a given protein across the three experimental replicates was calculated, and the log_2_-fold change in the average MASIC value for the test population relative to the ML control population was determined. Then, the log_2_-fold change for a given protein was compared to the log_2_-fold changes for all other proteins using a Z-score analysis. Log_2_-fold changes that were 2 standard deviations above or below the mean log_2_-fold change for the test population relative to the control population (Z-score < –2 or > 2) and for which the average MASIC value in both the test and the control population was greater than the limit of quantitation (LOQ) were considered significantly different. The full list of significant changes in protein abundance is available in Table S5. Significant differences between the three populations at a given time are represented as yellow or blue tiles in figures.

### Comparative analysis of expression changes

For comparative analyses of differentially expressed genes across the three populations, across time, and across the mRNA and protein levels, modified UpSet plots^137^ were generated using the R package *ComplexHeatmap* (v2.15.3)^138^.

### Significant differences in the lipidome

Significant differences in MGDG, DGDG, SQDG, and PG were assessed using a two-way mixed ANOVA using the R package *rstatix* (v0.7.2): normality was assessed using a Shapiro-Wilk test, and homogeneity of variance was assessed using a Levene test. Significant effects in accumulation of a given lipid class were classified as those which passed each test and had a *p* < 0.05, while significant effects in accumulation of specific individual lipid species within each lipid class were classified as those which passed each test and had a Bonferoni *p-adj.* < 0.05.

### Patterns and clustering of ‘omics data

Patterns of gene expression are represented as Z-scores of mean FPKMs for mRNAs and Z-scores of mean MASIC values for proteins across the 3 experimental replicates. Minimum and maximum FPKM and MASIC values are also shown to demonstrate the magnitude of change over time and across the three populations.

Patterns of lipid content are represented as Z-scores of the mean peak intensity across the 3 experimental replicates. As similar patterns over light intensity and time were observed across lipid species of the same class (MGDG, DGDG, and SQDG) (Figure S5B), the mean Z-scores and normalized peak intensities were calculated for each lipid class. Mean Z-scores of normalized abundances are used to demonstrate the differences over time and across the three populations, while the minimum and maximum normalized peak intensities are shown to demonstrate the dynamic range for each lipid class.

Principal-component analysis (PCA) of the transcriptome, proteome, and lipidome were performed in the R package *PCAtools* (v2.16.0), where the lower 10% of variables were removed according to variance. For the lipidome, PCA was performed separately for the data collected in negative mode and the data collected in positive mode. PCA of the TMT proteomics data revealed that, as expected, the primary data grouping component was TMT plex membership. Yet, PCA of individual plexes indicated high similarity across replicates and the influence of time and light intensity (data not shown). Therefore, we presented the PCA of the mean protein abundances across the experimental replicates to average out the contribution of TMT plex membership on the data.

The *k*-means clustering analysis was conducted with the R package *stats* (v4.4.0) using the k-means function. First, mean transcript abundances (as FPKMs) and mean protein abundances (as MASIC values) from all nuclear genes were independently Z-score normalized, and then merged. Genes that were undetected in one or both datasets were excluded, leaving 9,821 genes. The k-means function was applied to the resulting data frame with centers = 16 and iter.max = 1000. Next, genes were arranged by cluster assignment and a heatmap was generated with the heatmaps.2 function from the R package *gplots* (v3.1.3.1).

### Rhythmicity analysis

Rhythmic mRNA and protein accumulation were characterized for the LL, ML, and HL populations independently with the R package *DiscoRhythm* (v1.14.0)^45^ using a Cosinor algorithm^44^, which can tolerate time course data with non-equidistant sampling times. FPKM or MASIC values for the 3 experimental replicates were double-plotted, and the major expected period length was set to 24 h. The program excluded data with missing or constant values, so rhythmicity was assessed for roughly 13–14 x10^3^ transcripts and roughly 9 x10^3^ proteins in each of the three populations; full results of the analysis are available as Table S3. Pearson correlation and PCA were used to confirm that none of the samples was considered an outlier. Periods were detected and fit using a Cosinor model to principal component scores. Oscillations were detected using the Cosinor method, and accumulation was considered “rhythmic” if the q-value was < 0.05. To compare the phase of mRNA accumulation to the phase of protein accumulation, only genes whose products were rhythmic at both the mRNA and protein level for a given light intensity were considered (1004 genes in LL, 2379 genes in ML, 1849 genes in HL). To compare the phase of expression in the LL or HL populations relative to ML control population, only genes whose products were rhythmic in both light intensities were considered (roughly 11–12 x10^3^ transcripts, roughly 1–2 x10^3^ proteins).

### Gene ontology enrichment analysis

Previously determined gene ontology (GO) term assignments for all genes in *C. reinhardtii* v6.1 genome were downloaded from phytozome.net. Next, each cluster of genes from the *k*-means clustering analysis (Figure 2G), and for the core HL-responsive gene expression changes (Figure 3C) were analyzed for enrichment of GO terms with the R package *clusterProfiler* (v4.12.0) by using the enricher function with pvalueCutoff = 0.05, pAdjustMethod = “BH”. A representative subset of GO terms significantly enriched in the *k*-means clusters is displayed, and the full results of the enrichment analysis are available as Table S4. All significantly enriched GO terms are displayed for the core HL-responsive gene expression changes.

## Data availability

RNA-Seq raw and analyzed data have been deposited at NCBI Gene Expression Omnibus (GEO) and are publicly available as of the date of publication. Raw proteomics data have been deposited at the ProteomeXchange platform MassIVE and are publicly available as of the date of publication. Raw lipidomics data have been deposited at the National Metabolomics Data Repository (NMDR) and are publicly available as of the date of publication.

## ACKNOWLEDGMENTS

We thank Daniela Strenkert, Stefan Schmollinger, and M. Águila Ruiz-Sola for invaluable guidance on this project and critical reading of the manuscript. We gratefully acknowledge Charles Perrino for technical assistance, as well as Reena Zalpuri and Danielle Jorgens at the UC Berkeley Electron Microscopy Lab for EM sample preparation. We thank Holly L. Aaron and Feather Ives at the Molecular Imaging Center, UC Berkeley Cancer Research Laboratory for the technical setup for Airyscan microscopy (supported by NIH 1S10OD025063). We also thank Setsuko Wakao for her generosity with resources and instrument time. This work was supported by The Gordon and Betty Moore Foundation Symbiosis in Aquatic Systems Initiative Investigator Award GBMF9203 (https://doi.org/10.37807/GBMF9203) to S.S.M. and by the US Department of Energy (DOE) Office of Science through the Photosynthetic Systems program in the Office of Basic Energy Sciences funding to K.K.N. and M.I.. Proteomics and lipidomics were performed on a project award from the Environmental Molecular Sciences Laboratory, a DOE Office of Science User Facility sponsored by the Biological and Environmental Research program under Contract No. DE-AC05-76RL01830. V.O. acknowledges support from the European Union’s Horizon 2020 Research and Innovation Programme under the Marie Skłodowska-Curie grant agreement No. 887992. K.K.N. is an investigator of the Howard Hughes Medical Institute.

## AUTHOR CONTRIBUTIONS

V.O., M.I., and S.S.M. conceived of the study and designed the experiments. S.D. and V.O. conducted the experiments. S.D., V.O., A.G.G, C.D.N., and M.I. prepared and processed samples for physiological measurements, RNA-Seq, and proteomics. S.D., V.O., and R.P. performed image analysis. S.D., S.D.G., and S.O.P. analyzed the transcriptomic and proteomic data. S.D. and K.B. analyzed the lipidomic data. M.S.L., K.K.N, and S.S.M provided guidance and support for the study. S.D., V.O., and M.I. wrote the manuscript with input from all authors. S.S.M. edited the manuscript.

## SUPPLEMENTAL INFORMATION

**Document S1.** Figures S1–S7.

**Table S1.** Sample metadata for the transcriptome, proteome, and lipidome; related to Figures 2, 5, S1, and S5.

**Table S2.** Genome-wide mRNA abundance (FPKM) and protein abundance (MASIC value) for LL-, ML-, and HL-acclimated *Chlamydomonas* populations across the diurnal cycle in three experimental replicates; related to Figures 2 and S1.

**Table S3.** DiscoRhythm analysis of gene expression patterns in the LL-, ML-, and HL-acclimated populations at the mRNA and protein level; related to Figures 2 and S1.

**Table S4.** Gene ontology enrichment of the 16 clusters of genes; related to Figures 2 and S1.

**Table S5.** List of all significant changes in mRNA and protein abundance across the three photoacclimated populations at each timepoint; related to Figure 3.

**Table S6.** Core HL-responsive gene expression changes; related to Figures 3 and S4.

**Table S7.** The lipidome (median-normalized peak intensity) for LL-, ML-, and HL-acclimated *Chlamydomonas* populations across the diurnal cycle in three experimental replicates; related to Figures 5 and S5.

