## Supplementary material for "Too dim, too bright, and just right: Systems analysis of the *Chlamydomonas* diurnal program upon acclimation to light stress and limitation": Document S1

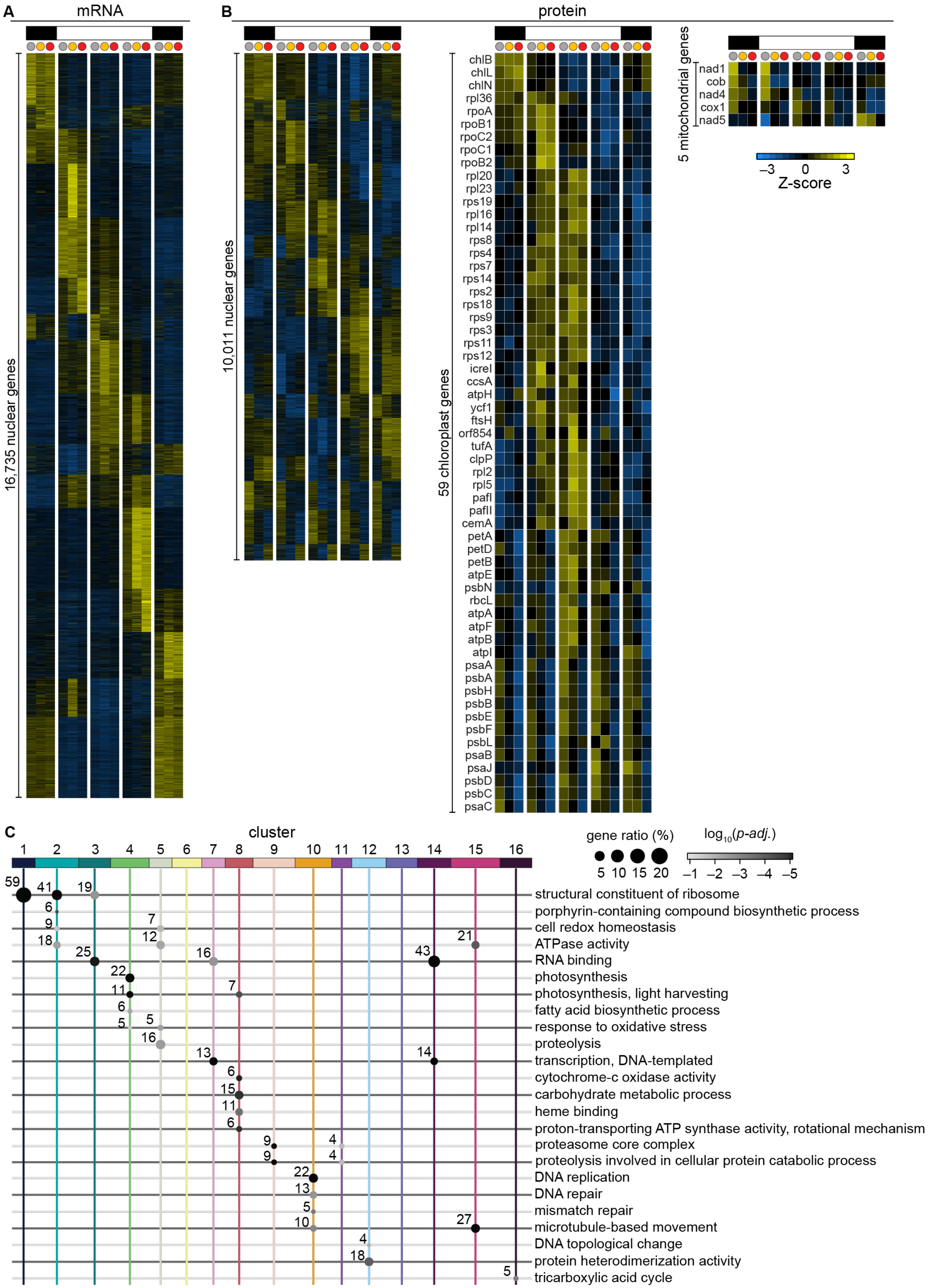

A
mRNA
16,735 nuclear genes
B
protein
10,011 nuclear genes
59 chloroplast genes
chlB
chlL
chlN
rpl36
rpoA
rpoB1
rpoC2
rpoC1
rpoB2
rpl20
rpl23
rps19
rpl16
rpl14
rps8
rps4
rps7
rps14
rps2
rps18
rps9
rps3
rps11
rps12
icreI
ccsA
atpH
ycf1
ftsH
orf854
tufA
clpP
rpl2
rpl5
pafI
pafII
cemA
petA
petD
petB
atpE
psbN
rbcL
atpA
atpF
atpB
atpI
psaA
psbA
psbH
psbB
psbE
psbF
psbL
psaB
psaJ
psbD
psbC
psaC
5 mitochondrial genes
nad1
cob
nad4
cox1
nad5
–3 0 3
Z-score
C
cluster
1 2 3 4 5 6 7 8 9 10 11 12 13 14 15 16
gene ratio (%)
5 10 15 20
log10(p-adj.)
–1 –2 –3 –4 –5
structural constituent of ribosome
porphyrin-containing compound biosynthetic process
cell redox homeostasis
ATPase activity
RNA binding
photosynthesis
photosynthesis, light harvesting
fatty acid biosynthetic process
response to oxidative stress
proteolysis
transcription, DNA-templated
cytochrome-c oxidase activity
carbohydrate metabolic process
heme binding
proton-transporting ATP synthase activity, rotational mechanism
proteasome core complex
proteolysis involved in cellular protein catabolic process
DNA replication
DNA repair
mismatch repair
microtubule-based movement
DNA topological change
protein heterodimerization activity
tricarboxylic acid cycle

**Figure S1: Genome-wide changes in mRNA and protein abundance over the LL, ML, and HL diurnal cycle; related to Figure 2.**

(A) Normalized abundance of 16,735 nucleus-encoded transcripts detected by RNA-Seq.

(B) Normalized abundance of 10,011 nucleus-encoded proteins, 59 chloroplast-encoded proteins, and 5 mitochondria-encoded proteins detected by TMT proteomics.

(C) Enrichment of selected GO terms in the 16 clusters in Figure 2G. Dot size indicates the proportion of genes in the cluster represented by the GO term, dot labels indicate the number of genes, and the shading indicates the $\log_{10}$*p-adj*. The full list of enriched GO terms is available as Table S4.

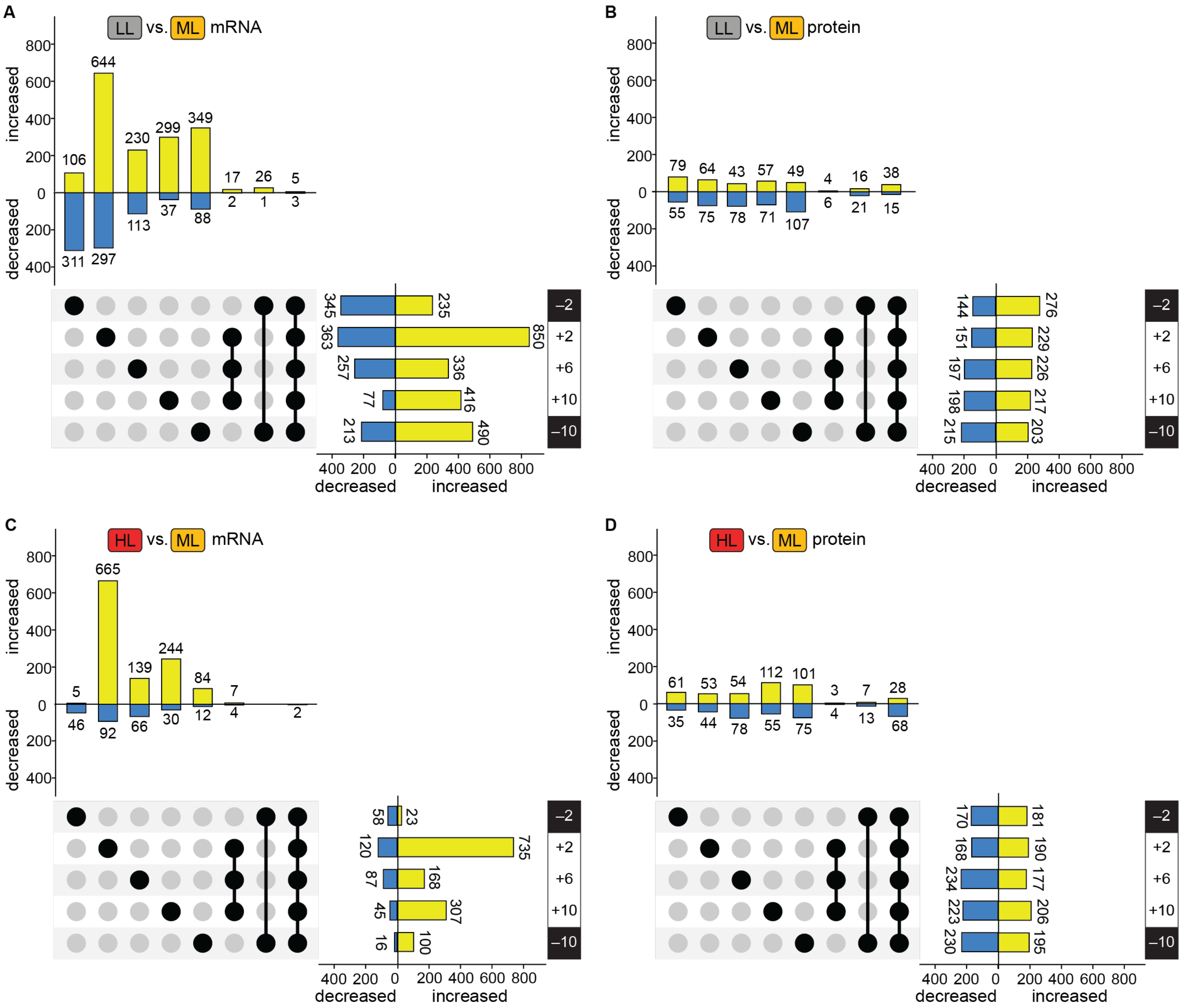
A
LL vs. ML mRNA
B
LL vs. ML protein
C
HL vs. ML mRNA
D
HL vs. ML protein
increased
decreased
–2
+2
+6
+10
–10
400 200 0 200 400 600 800
decreased
increased

**Figure S2: While most light-responsive changes in transcript abundance are specific to a particular time of day, protein abundance was often constitutively altered in the LL and HL populations; related to Figure 3.** Column bars represent the number of changes unique to each time, those common to the three light-phase timepoints, those common to the two dark-phase timepoints, and those common to all five timepoints (as indicated by the matrix below the columns). Row bars represent the total number of changes observed at each timepoint.

(A) Transient and constitutive changes in mRNA abundance in the LL population relative to the ML population.
(B) Transient and constitutive changes in protein abundance in the LL population relative to the ML population.
(C) Transient and constitutive changes in mRNA abundance in the HL population relative to the ML population.
(D) Transient and constitutive changes in protein abundance in the HL population relative to the ML population.

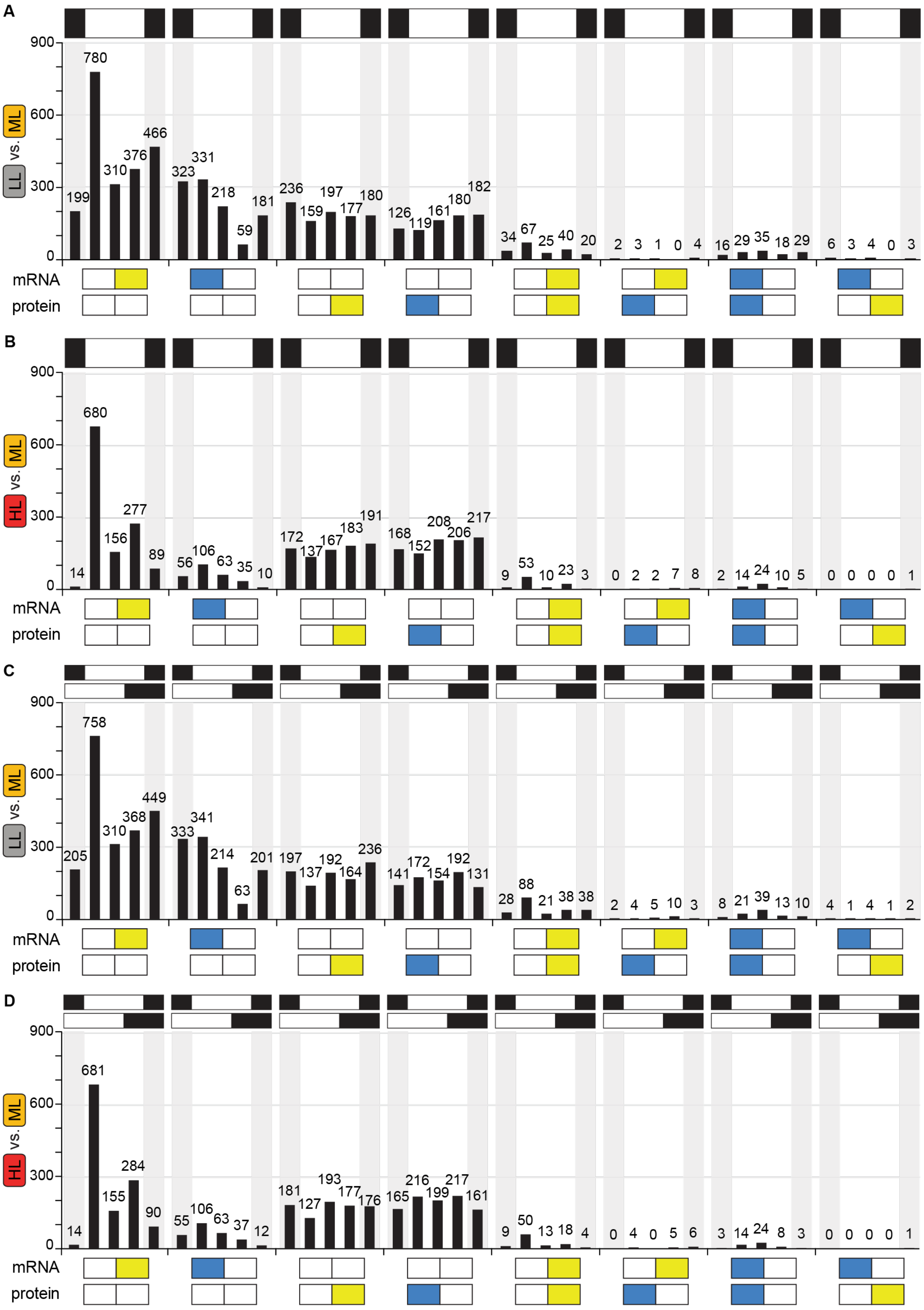
A
B
C
D
ML vs. LL
ML vs. HL
mRNA
protein
900
600
300
0

**Figure S3: Changes in gene expression are most often unique to the mRNA or protein level, even when accounting for a 4 h delay for protein accumulation; related to Figure 3.** The schematic x-axis represents the significant increases (yellow) and decreases (blue) that are unique or shared at mRNA and protein levels.

(A) Comparison of mRNA changes to protein changes for the LL population relative to the ML population.
(B) Comparison of mRNA changes to protein changes for the HL population relative to the ML population.
(C) Comparison of mRNA changes to protein changes 4 h later for the LL population relative to the ML population.
(D) Comparison of mRNA changes to protein changes 4 h later for the HL population relative to the ML population.

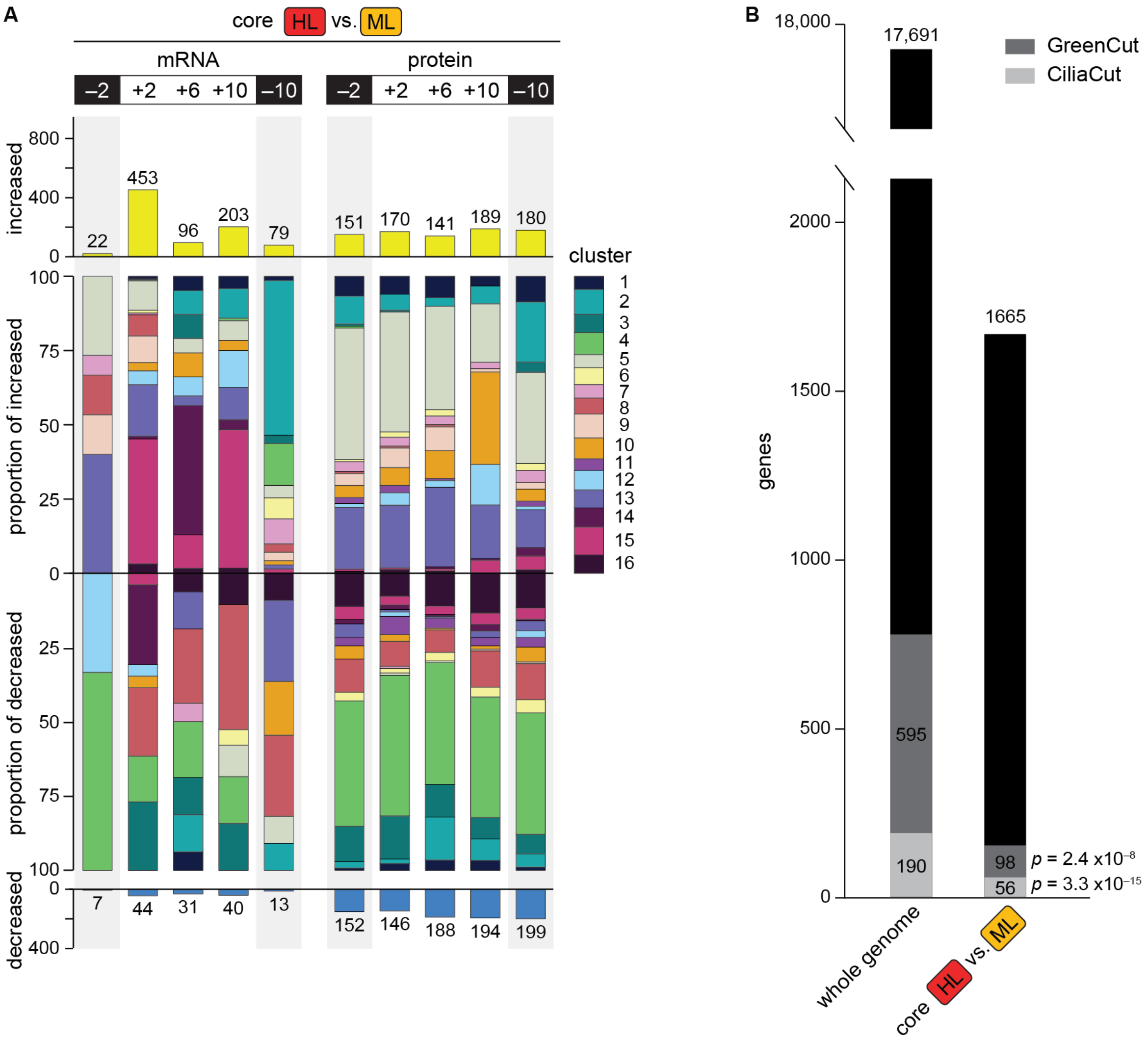


**Figure S4: Core HL-responsive gene expression changes; related to Figure 3.**

(A) The number of mRNAs significantly increased (top) and decreased (bottom) in abundance at each timepoint in the HL population relative to the ML population which are not shared with the LL population (core HL-responsive genes), and the proportion of those genes that belong to each cluster in Figure 2G (center); the key at right is scaled by the proportion of genes in the genome belonging to each cluster for comparison. A list of these changes is available as Table S6.

(B) The core HL-responsive genes are significantly enriched for GreenCut genes (genes retained across the green lineages and not in nonphotosynthetic organisms) and CiliaCut genes (genes retained in ciliated organisms and not in non-ciliated organisms)[55]. Enrichment *p* values are indicated.

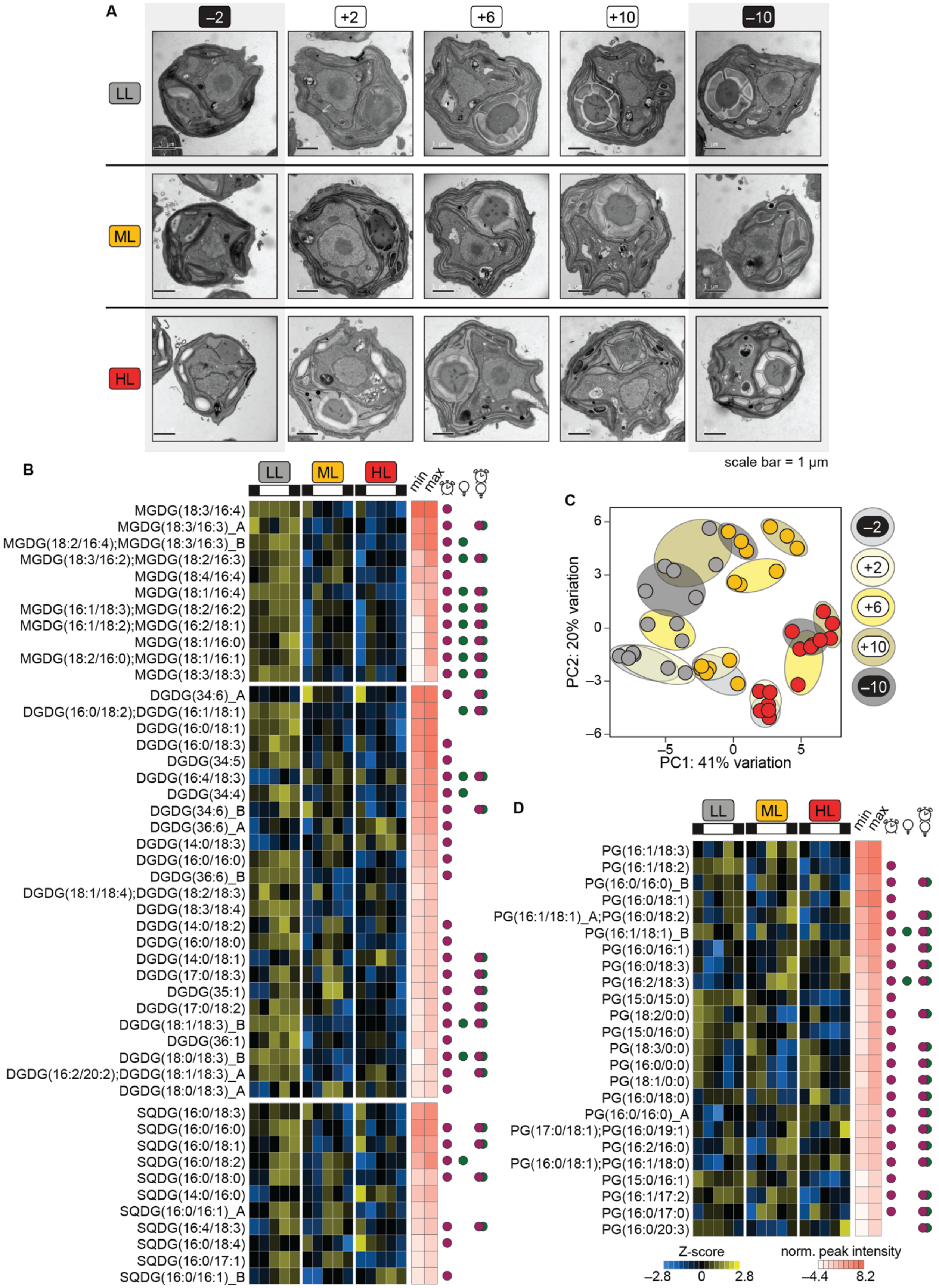
A
–2
+2
+6
+10
–10
LL
ML
HL
scale bar = 1 µm
B
LL
ML
HL
min
max
MGDG(18:3/16:4)
MGDG(18:3/16:3)_A
MGDG(18:2/16:4);MGDG(18:3/16:3)_B
MGDG(18:3/16:2);MGDG(18:2/16:3)
MGDG(18:4/16:4)
MGDG(18:1/16:4)
MGDG(16:1/18:3);MGDG(18:2/16:2)
MGDG(16:1/18:2);MGDG(16:2/18:1)
MGDG(18:1/16:0)
MGDG(18:2/16:0);MGDG(18:1/16:1)
MGDG(18:3/18:3)
DGDG(34:6)_A
DGDG(16:0/18:2);DGDG(16:1/18:1)
DGDG(16:0/18:1)
DGDG(16:0/18:3)
DGDG(34:5)
DGDG(16:4/18:3)
DGDG(34:4)
DGDG(34:6)_B
DGDG(36:6)_A
DGDG(14:0/18:3)
DGDG(16:0/16:0)
DGDG(36:6)_B
DGDG(18:1/18:4);DGDG(18:2/18:3)
DGDG(18:3/18:4)
DGDG(14:0/18:2)
DGDG(16:0/18:0)
DGDG(14:0/18:1)
DGDG(17:0/18:3)
DGDG(35:1)
DGDG(17:0/18:2)
DGDG(18:1/18:3)_B
DGDG(36:1)
DGDG(18:0/18:3)_B
DGDG(16:2/20:2);DGDG(18:1/18:3)_A
DGDG(18:0/18:3)_A
SQDG(16:0/18:3)
SQDG(16:0/16:0)
SQDG(16:0/18:1)
SQDG(16:0/18:2)
SQDG(16:0/18:0)
SQDG(14:0/16:0)
SQDG(16:0/16:1)_A
SQDG(16:4/18:3)
SQDG(16:0/18:4)
SQDG(16:0/17:1)
SQDG(16:0/16:1)_B
C
PC2: 20% variation
PC1: 41% variation
6
3
0
–3
–6
–5
0
5
–2
+2
+6
+10
–10
D
LL
ML
HL
min
max
PG(16:1/18:3)
PG(16:1/18:2)
PG(16:0/16:0)_B
PG(16:0/18:1)
PG(16:1/18:1)_A;PG(16:0/18:2)
PG(16:1/18:1)_B
PG(16:0/16:1)
PG(16:0/18:3)
PG(16:2/18:3)
PG(15:0/15:0)
PG(18:2/0:0)
PG(15:0/16:0)
PG(18:3/0:0)
PG(16:0/0:0)
PG(18:1/0:0)
PG(16:0/18:0)
PG(16:0/16:0)_A
PG(17:0/18:1);PG(16:0/19:1)
PG(16:2/16:0)
PG(16:0/18:1);PG(16:1/18:0)
PG(15:0/16:1)
PG(16:1/17:2)
PG(16:0/17:0)
PG(16:0/20:3)
Z-score
–2.8
0
2.8
norm. peak intensity
–4.4
8.2

**Figure S5: Diurnal photoacclimation leads to stable changes in thylakoid membranes and individual chloroplast lipid species that persist in the dark phase; related to Figure 5.**

(A) Representative electron micrographs of whole *Chlamydomonas* cells from the 15 cellular states assayed in this study.

(B) Changes in individual MGDG, DGDG, and SQDG lipid species. Circles at right indicate a significant effect of time, light intensity, or the interaction between the two by two-way mixed ANOVA (Bonferroni *p-adj* < 0.05).

(C) PCA of lipids detected by LC-ESI-MS/MS in negative ionization mode (PG, PA, PE, and PI species); ellipses designate time of day.

(D) Changes in individual PG lipid species. Circles at right indicate a significant effect of time, light intensity, or the interaction between the two by two-way mixed ANOVA (Bonferroni *p-adj* < 0.05).

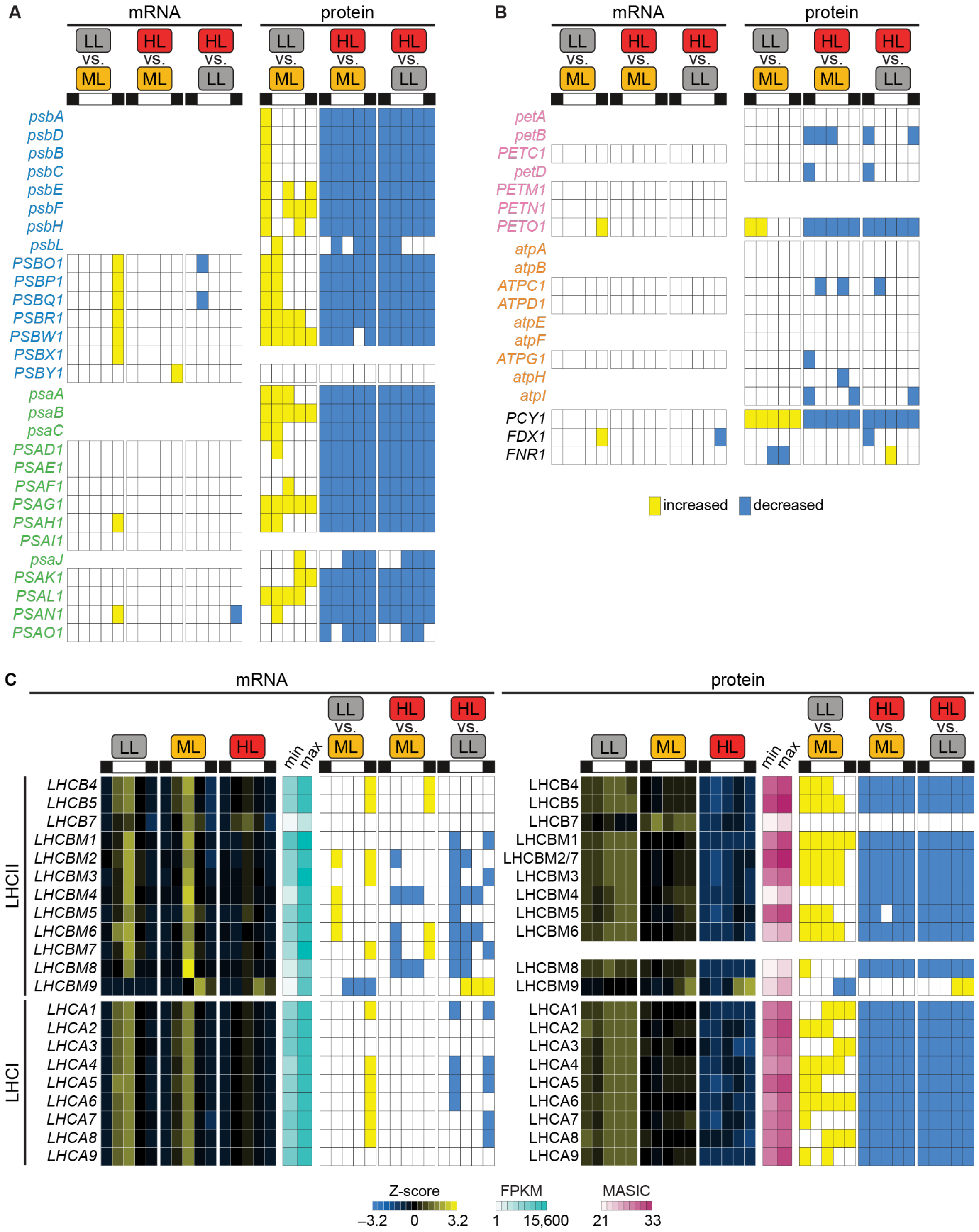


**Figure S6: Diurnal photoacclimation results in stable changes to LHC and photosystem protein abundances; related to Figure 6.**

(A) Significant changes in photosystem mRNA and protein abundances across the three populations. Gene names are colored by complex as in Figure 6.

(B) Significant changes in mRNA and protein abundances for other components of the photosynthetic electron transfer chain across the three populations. Gene names are colored by complex as in Figure 6.

(C) Changes in the expression of LHCs. Individual LHC transcripts and proteins show similar patterns over time in the three populations, except for LHCBM9 and LHCB7.

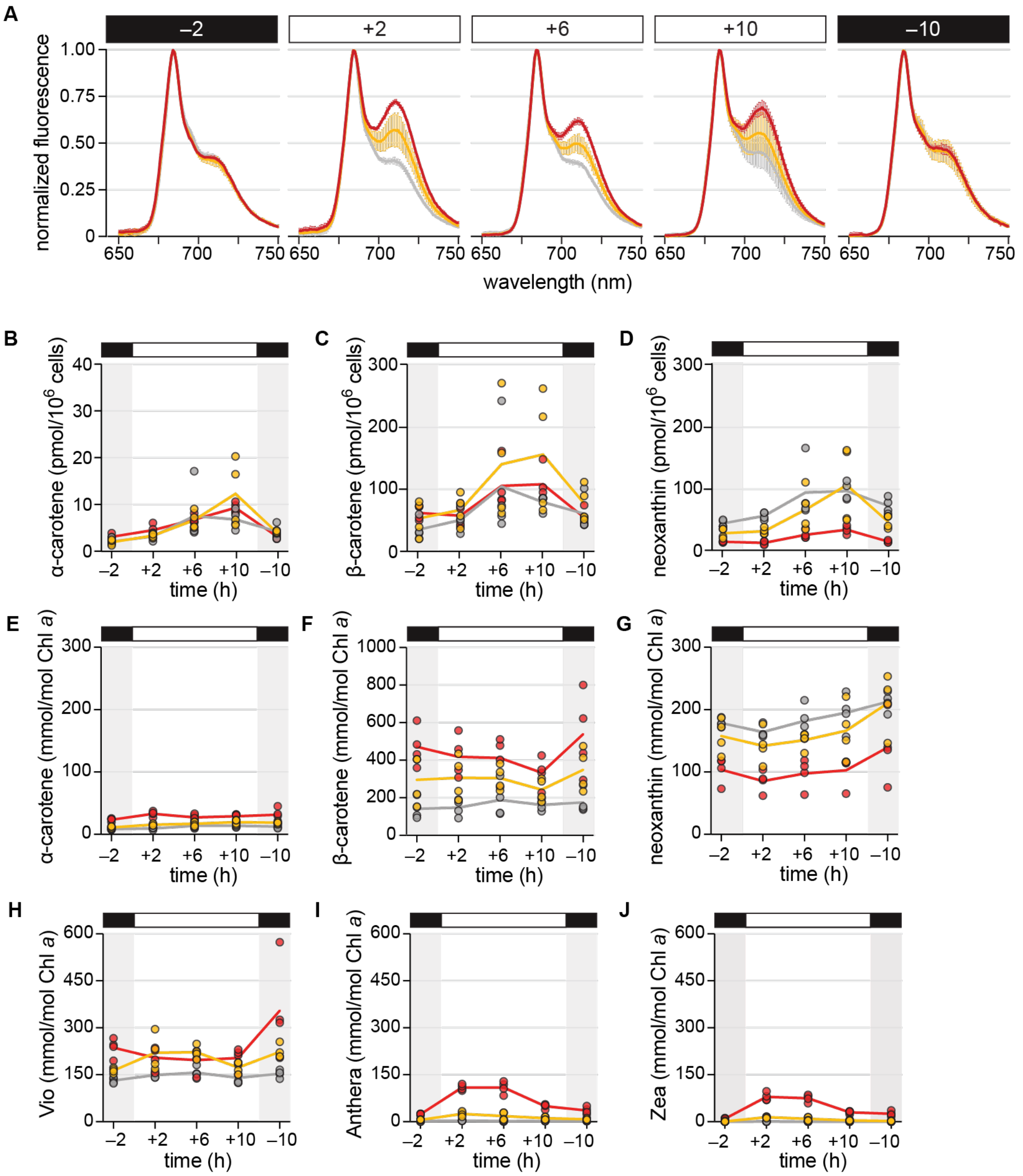


**Figure S7: The *Chlamydomonas* antenna and cellular pigment profile is dynamic over the diurnal cycle and in response to light intensity; related to Figure 7.**

(A) 77 K fluorescence emission spectra are altered during the light phase in the ML and HL populations. Data are represented as the mean of 4 experimental replicates (n = 4) with error bars representing the standard deviation.

(B) Cellular α-carotene concentration.

(C) Cellular β-carotene concentration.

(D) Cellular neoxanthin concentration.

(E) Changes in α-carotene relative to Chl *a*.

(F) Changes in β-carotene relative to Chl *a*.

(G) Changes in neoxanthin relative to Chl *a*.

(H) Changes in Vio relative to Chl *a*.

(I) Changes in Anthera relative to Chl *a*.

(J) Changes in Zea relative to Chl *a*.
